# Yellow Fever Virus Interactomes Reveal Common and Divergent Strategies of Replication and Evolution for Mosquito-borne Flaviviruses

**DOI:** 10.1101/2025.06.14.659623

**Authors:** Matthew W. Kenaston, Liubov Cherkashchenko, Chase L. S. Skawinski, Adam T. Fishburn, Vardhan Peddamallu, Cole J. Florio, Avery E. Robertson, Tamanash Bhattacharya, Janet M. Young, Harmit S. Malik, Priya S. Shah

## Abstract

Pathogenic mosquito-borne flaviviruses infect mosquito and human hosts, relying on host protein interactions to replicate, evade immunity, and mediate pathogenesis. Prior proteomic studies mapped such interactions for some flaviviruses, but yellow fever virus (YFV)—a pathogen of resurgent concern—remains understudied. Here, we map YFV interactomes in human and mosquito cells to identify interactions common among divergent flaviviruses or unique to YFV. Functional assays reveal a previously unrecognized YFV restriction factor: RBBP6 inhibits YFV genome replication by interacting with the viral polymerase NS5. We enhance the identification of dual-host interactions using structural modeling and holistic network integration. Extending our holistic approach to other flavivirus interactomes, we distinguish conserved mechanisms of host targeting from those unique to YFV. Contrary to expectations that conserved viral proteins lead to conserved protein interactions, we find that Capsid, a divergent structural protein, shares more host interactions than NS5, a conserved enzyme. Integrating proteomics with complementary analyses defines new principles of host-targeting strategies across flavivirus and host evolution, offering a versatile resource for navigating the complex landscape of flavivirus biology.

## Introduction

Mosquito-borne orthoflaviviruses (commonly known as flaviviruses) pose a serious and increasing threat to public health worldwide with their continued re-emergence in the past decade. These positive-sense RNA viruses, which includes dengue virus (DENV), Zika virus (ZIKV), West Nile virus (WNV), and yellow fever virus (YFV), contributes to hundreds of millions of human infections worldwide every year^1^. Due to ongoing climate change, the geographical range of common vectors, such as the *Aedes aegypti* mosquito, is rapidly expanding^2,3^. Human urbanization and encroachment have further exacerbated this trend^4,5^. And yet, there are only limited vaccine strategies and no approved antivirals for flaviviruses^6^. Thus, it is critical to unravel the molecular mechanisms governing flavivirus replication.

Identifying virus-host protein-protein interactions (PPIs) is crucial for developing both virus- and host-targeted therapies^7^. Flaviviruses have small RNA genomes that encode a single polyprotein, which is subsequently proteolytically processed into 10 viral proteins. Three of these proteins—Capsid, prM (membrane precursor), and Envelope—are structural components of the virion, while the remaining non-structural (NS) proteins facilitate replication, translation, assembly, and viral egress. These viral proteins use physical interactions with host proteins to hijack cellular machinery essential for viral replication, immune evasion, and pathogenesis^8^. Previous studies have applied proteomics-based approaches to identify virus-host PPIs across vertebrate and invertebrate host cells^9,10^, offering new strategies for vector control and therapeutic targets.

Due to their severe disease burdens or alarming pathogenesis, flaviviruses like DENV and ZIKV have been the focus of many previous studies^11^. In contrast, YFV has yet to be studied using contemporary proteomic techniques. This is partly because a robust live attenuated vaccine^12^ can provide long-term immunity^13^, reducing the perceived threat of YFV. However, limited vaccine availability and efficacy combined with unexpected increases in mosquito vector competency have led to repeated YFV outbreaks in the last decade^14–16^. YFV was estimated to cause 109,000 severe infections and 51,000 deaths in 2018 alone^17^. More recently, a Pan-American Health Organization alert in February 2025 reported an alarming increase in YFV cases in the Americas, prompting urgent recommendations for enhanced protection^18^. Recent surveillance also suggests YFV range is expanding^19^. Considering this resurgence, understanding the molecular mechanisms of YFV replication and pathogenesis has regained urgency and relevance.

YFV is an important virus to consider from an evolutionary perspective. YFV is phylogenetically distinct from DENV, ZIKV, and WNV^20,21^, flaviviruses whose host interactions have been studied in depth^22–24^. DENV, ZIKV, and WNV are more closely related to each other than to YFV. However, DENV, ZIKV, and YFV can all use *A. aegypti* as a vector and use primates as vertebrate reservoirs, whereas WNV employs *Culex* mosquitoes as vectors and birds as amplifying vertebrate reservoirs^1^. Analyzing YFV-host PPIs thus represents a powerful opportunity to understand the biology of an understudied and biomedically relevant flavivirus while also revealing unique and broadly conserved features of flavivirus biology across different flavivirus-host PPI datasets.

Here, we conduct comprehensive proteomics analyses to construct YFV-host PPI networks in both human and mosquito (*A. aegypti*) cells. Using a targeted CRISPRi knockdown screen of YFV-interacting human proteins, we identify the E3 ubiquitin ligase RBBP6 (retinoblastoma binding protein 6) as a novel YFV restriction factor. We integrate computational approaches, including evolutionary conservation metrics, AI-based structural modeling, and high-dimensional network-based analyses, to systematically evaluate the breadth of flavivirus-host PPIs across hosts and viruses. Our studies improve the identification of shared interactions across divergent hosts and reveal general flaviviral strategies as well as YFV-specific interactions that reflect YFV’s divergence from other flaviviruses. Finally, by integrating PPI studies across four flaviviruses, we discover a counterintuitive trend regarding the conservation of virus-host PPIs. The conservation of flavivirus proteins is inversely correlated with the conservation of their host interactomes, raising interesting questions regarding the speed at which some highly conserved flavivirus proteins can evolve new protein interactions. By integrating multiple proteomic datasets with orthogonal experimental and computational approaches, we create a multidimensional resource for understanding host interactions, replication strategies, and evolution of mosquito-borne flaviviruses.

## Results

### A YFV-Human Interactome

To map the YFV-human PPI network, we employed a proteomics-based approach using affinity purification-mass spectrometry (AP-MS). We first cloned protein-coding sequences from the Asibi strain of YFV (YFV-Asibi) with C-terminal 2xStrep affinity tags. Following transient expression in HEK293T cells, we performed affinity purification and confirmed the expression and purification of both viral baits and co-purified host proteins (Figures 1A and S1A). We used LC-MS/MS to analyze eluted proteins. Replicate samples associated with specific viral baits showed strong correlations with each other but not with samples from other viral baits, indicating reproducibility and specificity (Figure S1B). To score these interactions, we applied a tiered computational approach, integrating Mass Spectrometry Interaction STatistics (MiST) and Significance Analysis of INTeractome (SAINT) (Table S1). We identified 309 high-confidence PPIs (HC-PPIs) spanning 10 YFV baits (Figure 1B). We only failed to identify HC-PPIs for prM protein, which also had few interactions in past studies of other flaviviruses using a similar approach^22,23^.

**Figure 1.**
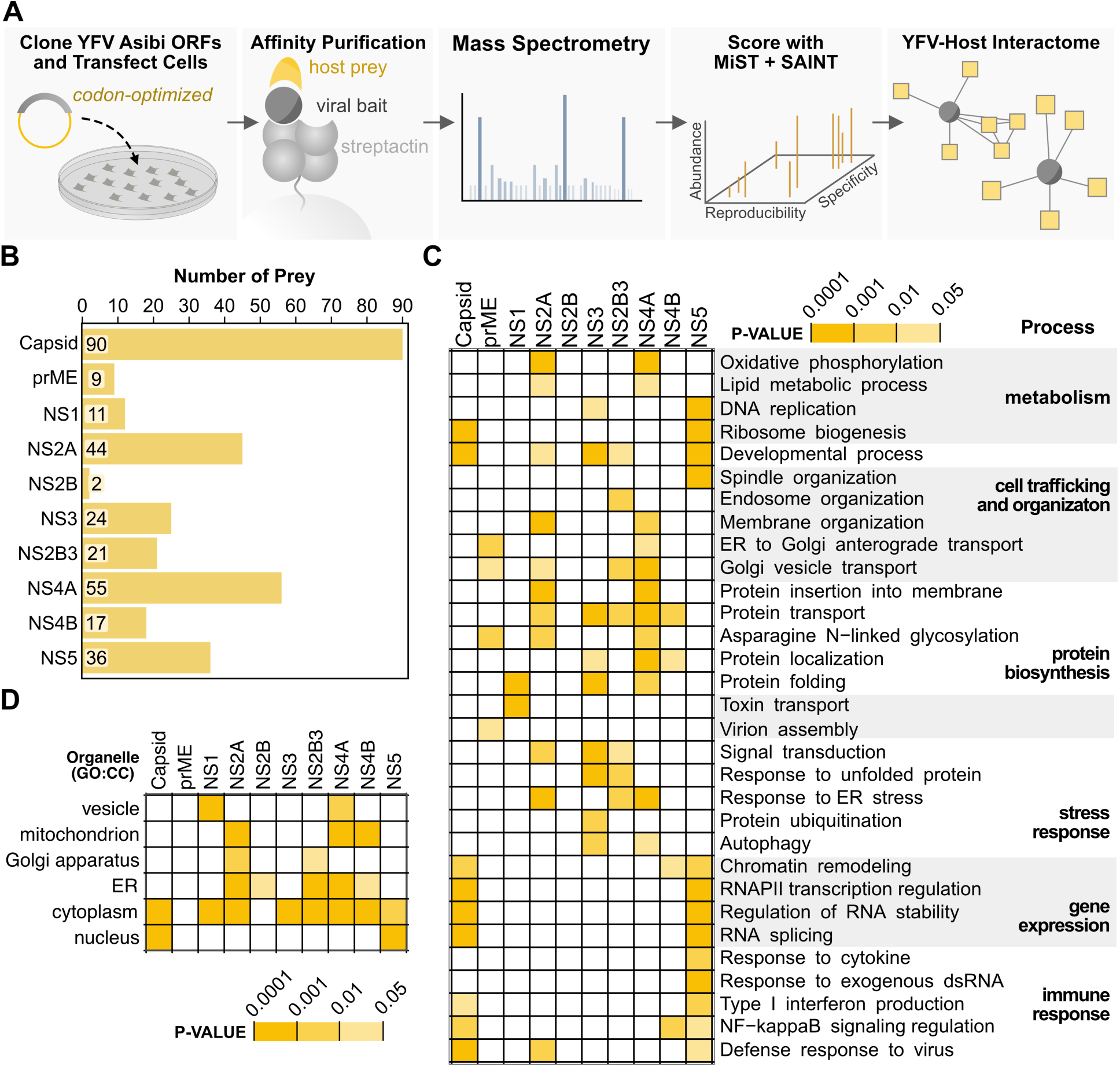
YFV-human proteomics identifies virus-host PPIs related to flavivirus biology. (A) Schematic representation of experimental AP-MS pipeline and proteomic scoring. (B) Virus-host interactions were scored with both MiST and SAINT to produce the plotted number of HC-PPIs per bait. (C-D) Enrichment analysis was performed on the set of HC-PPIs for each viral bait. The heatmaps summarize the degree of enrichment (adjusted p-value < 0.05) for (C) GO: Biological Processes and Reactome pathways relevant to flavivirus replication, and (D) GO:cellular compartment. The full enrichment results are available in Table S1. Viral baits are ordered according to their position in the viral polyprotein.

To broadly assess the functional relevance of YFV-human PPIs, we performed enrichment analyses on the HC-PPIs associated with each viral protein using Gene Ontology: Biological Process (GO:BP) and Reactome pathway analyses (Figure 1C and Table S1). These analyses revealed the broad targeting of host processes previously shown to be critical for flavivirus replication and pathogenesis^9^, including RNA metabolism^25–27^, chromatin modification and remodeling^22,28^, ER stress response^29^, and cell trafficking and organization^30,31^. NS5 and Capsid exhibited strong enrichment for nuclear processes, such as RNA splicing, antiviral defense, and ribosome biogenesis, consistent with their previously characterized nuclear functions^28,32–35^. NS4A and NS2A primarily targeted pathways related to membrane dynamics, including protein insertion into membranes and ER organization^36,37^. NS3 and NS2B3 showed distinct yet complementary enrichment patterns, with NS2B3 favoring endosomal pathways and NS3 interacting with ubiquitination and protein folding machinery. We found that prME, the protein complex of the Envelope (E) with its chaperone prM, is uniquely associated with pathways involved in virion assembly, potentially reflecting its role in flavivirus particle maturation. The subcellular localization of host proteins involved in HC-PPIs also correlated well with our observed subcellular localizations of the respective flavivirus proteins (Figures 1D and S2). To contextualize our investigation and provide visualization, we constructed a network of YFV-human HC-PPIs annotated with GO:BP and CORUM protein complexes^38^ (Figure 2). This network provides a comprehensive resource for understanding YFV-human interactions.

**Figure 2.**
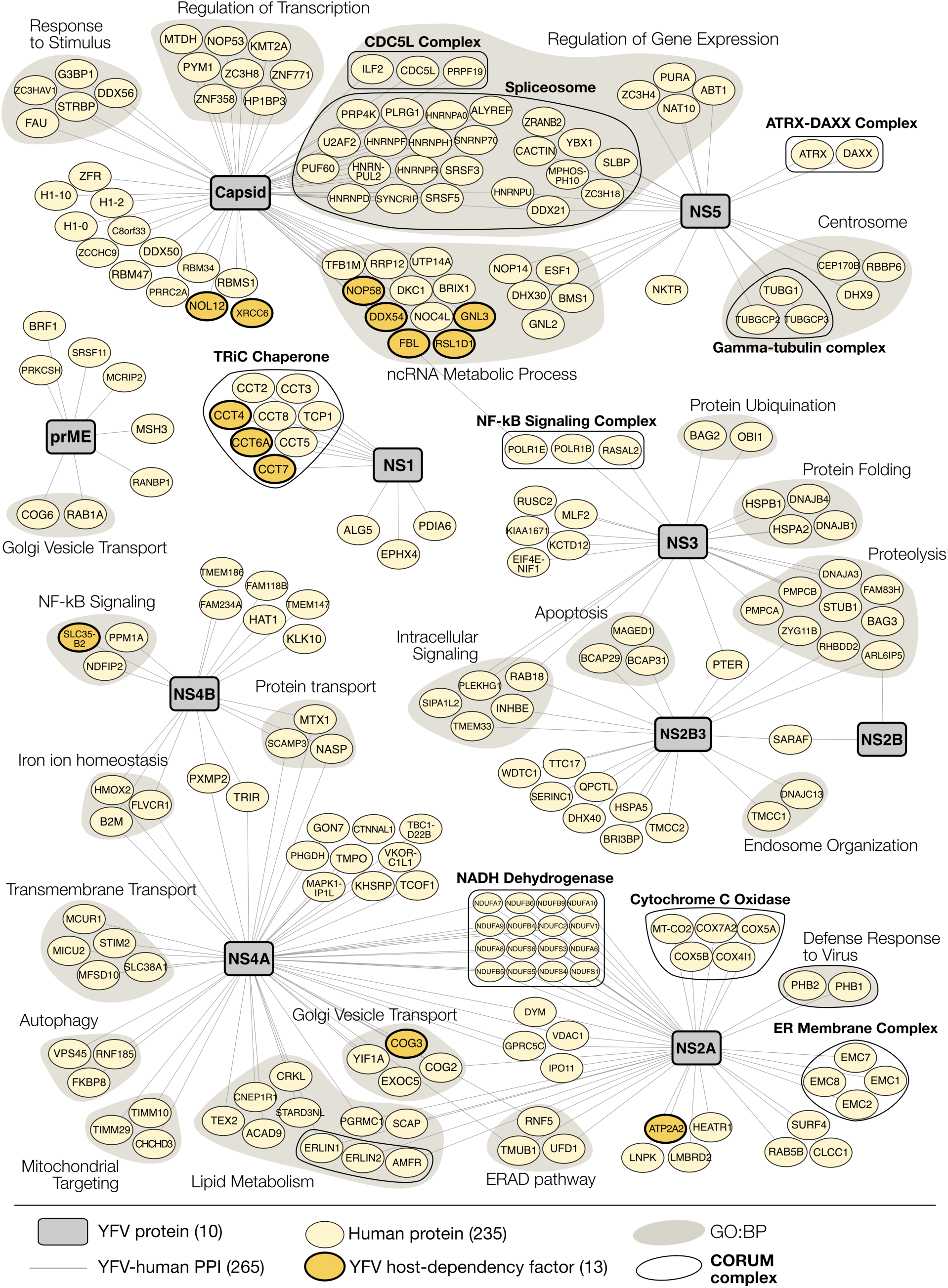
YFV-human PPI network. YFV-Human HC-PPIs were identified after a tiered scoring approach using both MiST (M > 0.67 | M > 0.60 & protein complex member) and SAINT (score > 0.95 & BFDR ≤ 0.05). Viral baits (grey rectangles) are mapped to human proteins (light yellow ovals). Human proteins are annotated as host-dependency factors from a contemporary YFV CRISPR screen (gold with bolded borders). Additionally, human proteins were grouped by GO:BP (grey underlay) and CORUM protein complex (black outline). Full scoring results are available in Table S1. Ribosomal proteins (42) were not included for visual clarity.

### RBBP6 is a Novel Inhibitor of YFV Genome Replication

Genome-wide screening approaches for host dependency factors have revealed many important details of flavivirus replication^35,37,39–42^. We compared our YFV-human HC-PPIs with a recent YFV-focused genome-wide CRISPR screen that identified host dependency factors in human cells^35^. We identified a significant overlap (p = 2.8x 10^-14^) of 19 host factors among our 277 unique prey and the top 1% of YFV host dependency factors (Figure 2). This suggests our HC-PPIs are enriched for host proteins involved in YFV replication. However, this is likely an underestimate, since many flavivirus host factors cannot be studied using pooled genome-wide approaches due to their inherent limitations, such as the selection criteria and the targeting of essential genes. Proteomics-based interactomes offer a powerful complementary approach to genome-wide screens, providing a targeted set of host genes to assess the impacts on viral replication.

To further dissect the functional consequences of promising YFV-host interactions, we performed a targeted CRISPR interference (CRISPRi) screen of 18 host proteins from our YFV-human HC-PPI network. We modified our experimental setup in two ways compared to our proteomics assays. First, we used the 17D strain of YFV (YFV-17D), a biosafety level 2 strain that is more amenable to infection-based screening. Despite serial passaging, only 32 amino acids differentiate the entire polyproteins of YFV-17D and YFV-Asibi^43^. Thus, YFV-17D retains a high degree of sequence similarity to YFV-Asibi, which was used to generate our proteomic data. Second, since the liver is a major site of YFV replication^44–46^, we carried out our screen in hepatocyte-derived Huh7 cells instead of the HEK293T cells used for proteomics.

Using transient CRISPRi knockdown in Huh7-dCas9 cells, we assessed the effect of host protein depletion on YFV-17D titers. We individually targeted each host gene with two unique guide RNAs (gRNAs); most gRNAs achieved efficient knockdown (>70%). We proceeded with any gRNAs that reached at least 50% knockdown relative to the non-targeting negative control (ncgRNA) (Figure 3A). Knockdown cells were then infected with YFV-17D at a multiplicity of infection (MOI) of 0.1, and infectious viral titers were quantified by plaque assay at 48 hours post-infection (hpi). We observed strong reproducibility across three biological replicates, with a Pearson’s correlation coefficient (R) ranging from 0.62 to 0.80. Overall, 10 of 30 tested gRNAs (33%), targeting 7 unique genes, significantly influenced YFV titers (Figure 3B and Table S2, |robust Z-score| > 2). Our findings are consistent with previous studies examining host factors involved in flavivirus replication^23^. Viral replication decreased in two knockdowns targeting two unique host genes (robust Z-score < −2), suggesting these host genes encode dependency proteins that perform proviral functions. In contrast, 8 knockdowns targeting 5 unique host genes resulted in increased viral replication (robust Z-score > 2); these host genes likely represent antiviral factors that restrict viral replication.

**Figure 3.**
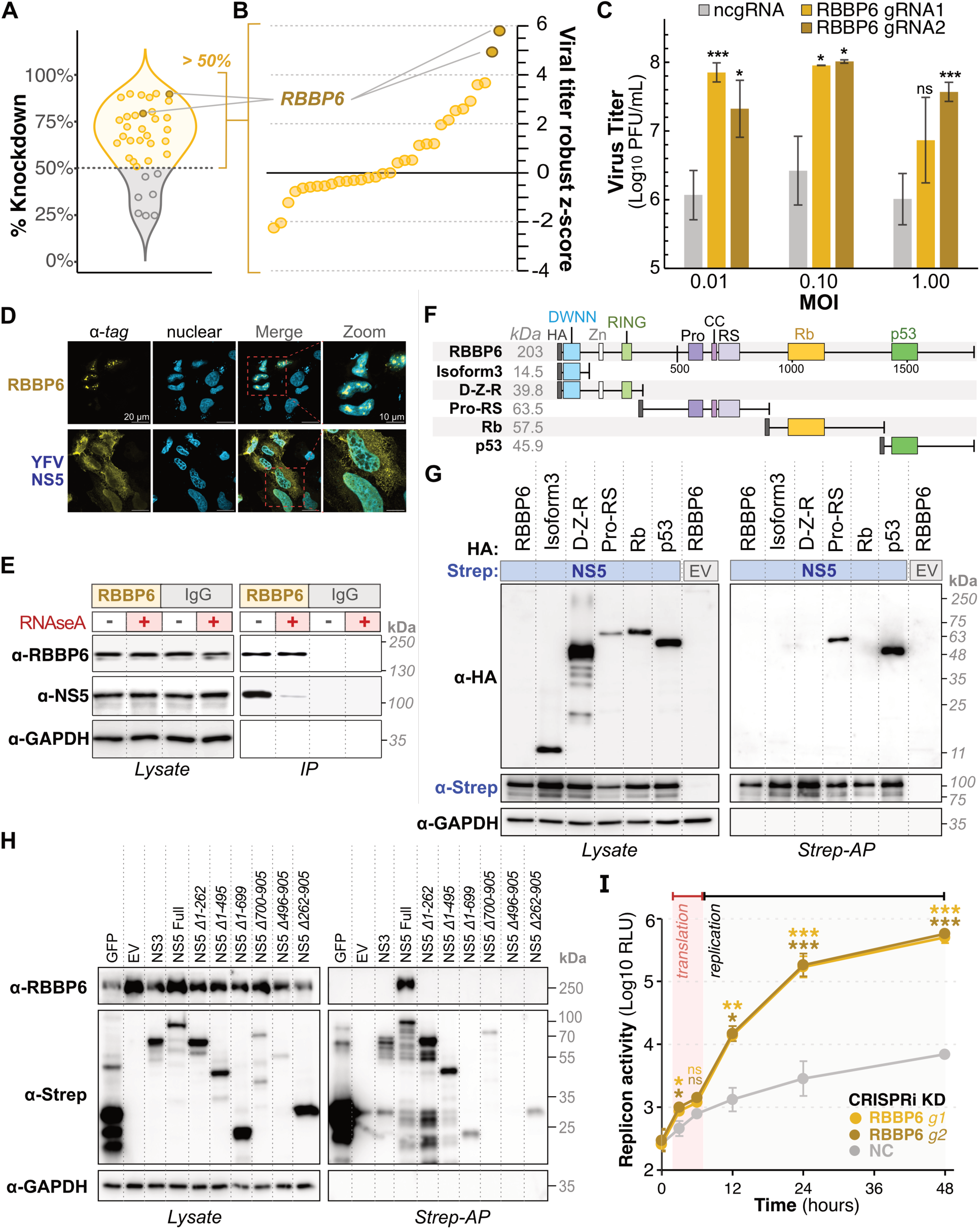
RBBP6 inhibits YFV genome replication through an RNA-dependent interaction with full-length NS5. (A) Violin plot showing the percent knockdown efficiency of all tested gRNAs in the CRISPRi screen. gRNAs achieving achieved >50% knockdown are shown in yellow. RBBP6-targeting gRNAs are shown in bronze. (B) Robust Z-score distribution of viral titers from the CRISPRi screen for gRNAs achieving >50% knockdown (panel A, yellow), identifying RBBP6 as a top restriction factor (bronze). Full results are available in Table S2. (C) YFV-17D replication in Huh7 cells with RBBP6 knockdown. Plaque assay quantification of viral titers at 48 hours post-infection (hpi) across a range of multiplicities of infection (MOI = 0.01, 0.10, 1.00). Data represent mean ± SD of three biological replicates, with p < 0.005 (***), p < 0.01 (**), and p < 0.05 (*) by one-sided t-test. (D) Confocal immunofluorescence microscopy of RBBP6 and YFV NS5 in transfected HeLa cells. 2xStrep-tag for NS5 and HA-tag for RBBP6 (yellow). Hoechst staining for nucleus (cyan). Scale bars: 20 µm (main panels), 10 µm (zoom). (E) Western blot for co-IP of NS5 with endogenous RBBP6 or IgG in YFV-infected cells with/without RNaseA treatment. GAPDH is a loading control. (F) Schematic of RBBP6 domain architecture and truncation constructs. (G) Western blot for whole cell lysate and Strep-AP of YFV 2xStrep-tagged NS5 co-transfected with HA-tagged RBBP6 constructs. EV represents an empty vector for negative control. (H) Western blot for whole cell lysate and Strep-AP of YFV 2xStrep-tagged NS5 domain truncations. NS3 and EV are negative controls. (I) *Renilla* luciferase activity of YFV replicon in RBBP6 and control CRISPRi knockdown Huh7 cells. Data represent mean ± SD of three biological replicates, with p < 0.005 (***), p < 0.01 (**), and p < 0.05 (*) by one-sided t-test.

Most striking among the putative restriction factors was RBBP6. A >75% reduction in RBBP6 expression (Figure 3A) resulted in up to 60-fold increase in YFV titers for two different gRNAs across multiple MOIs (Figure 3C). This suggests that RBBP6 potently restricts YFV replication. RBBP6 is an E3 ubiquitin ligase that regulates RNA 3’ end processing, the cell cycle, and embryonic development^47–49^. Our PPI analyses revealed that RBBP6 interacts with YFV NS5 (Figure 2), which encodes the RNA-dependent RNA polymerase (RdRP) and methyltransferase (MTase). Both YFV NS5 and RBBP6 can localize to the nucleus (Figure 3D). To validate this interaction during infection, we performed endogenous RBBP6 immunoprecipitation (IP) in YFV-17D-infected Huh7 cells. Western blotting revealed that NS5 co-precipitated with RBBP6 (Figure 3E). Additionally, treating IP samples with RNaseA substantially decreased the amount of NS5 that could be co-immunoprecipitated with RBBP6, indicating the interaction is stabilized by RNA (Figure 3E).

Next, we used domain deletions to map the domains of RBBP6 responsible for this interaction (Figure 3F). Although full-length RBBP6 was poorly expressed, smaller RBBP6 domains were expressed well (Figure 3G). Co-IP analyses revealed that both the Proline-Rich (Pro-RS) and p53-binding domains were independently sufficient for interaction with NS5 (Figure 3G). Confocal immunofluorescence microscopy revealed clear nuclear localization for the Pro-RS domain; however, the p53-binding domain had a cytoplasmic localization (Figure S3A-C). We propose that YFV NS5 interaction with the p53 domain of RBBP6 may be facilitated by NS5 nucleocytoplasmic shuttling (Figures S2A and S3D-F).

We also mapped the NS5 domains required for interaction with RBBP6 by co-IP assays. While full-length NS5 interacts with RBBP6, neither the MTase (residues 1 to 261) nor RdRP (residues 262 to 909) domain was sufficient to pull down RBBP6 (Figure 3H, Figure S3D). No YFV NS5 truncations were excluded from the nucleus, suggesting that their lack of interaction with RBBP6 was unrelated to subcellular mislocalization (Figure S3D-F). Our findings suggest that YFV NS5 requires an intact conformation or the presence of multiple domains to bind RBBP6.

NS5 encodes for the viral RdRP and MTase, and is therefore essential for viral RNA synthesis. We tested whether RBBP6 depletion enhanced YFV genome replication using a replicon encoding a luciferase reporter^50^. The flavivirus life cycle is characterized by an initial phase of translation, in which little to no increase in reporter activity is observed, followed by a switch to replication, during which reporter activity increases dramatically. Depletion of RBBP6 resulted in minor but significant increases in reporter activity during the early translation phase and a more pronounced and sustained ∼10-to 70-fold increase in reporter activity during the replication phase (Figure 3I). Together, our data support a model in which RBBP6 acts as a YFV restriction factor by inhibiting YFV genome replication through a complex RNA-mediated interaction with NS5.

Positive selection is often observed in other antiviral restriction factors at their interface with viral targets due to an arms race for binding affinity or evasion from binding^51–54^. However, analysis of 28 primate *RBBP6* sequences revealed no significant evidence of positive selection (Table S3). This lack of positive selection might reflect a lack of significant coevolution between flaviviruses and primate *RBBP6* or constraints on *RBBP6* to maintain its essential host functions.

### A Structure-based and Holistic Approach to Identify Interacting Homologs

Mosquito-borne flaviviruses must replicate in both vertebrate and invertebrate hosts for their transmission. They do so by hijacking host processes across evolutionarily distinct cellular environments^37,40^. To identify how YFV proteins commandeer mosquito cells, we constructed a YFV-mosquito interactome following the same steps as our YFV-human network, except that we expressed YFV proteins in *A. aegypti* Aag2-AF5 cells^55^ (Figure S1C). We identified YFV-mosquito HC-PPIs using the same AP-MS and computational scoring pipeline previously employed for human cells, ensuring comparability across datasets (Figures 1A and S1D, and Table S4). Despite lower detection sensitivity in mosquito cells, we identified robust interactions for most YFV proteins (Figure 4A). The only exceptions were prM and prME, which failed to express in Aag2-AF5 cells (Figure S1C). Furthermore, although NS1 and NS2B were expressed well, none of their interactions met our scoring threshold. In total, we identified 150 YFV-mosquito HC-PPIs across 7 viral baits (Figure 4A).

**Figure 4.**
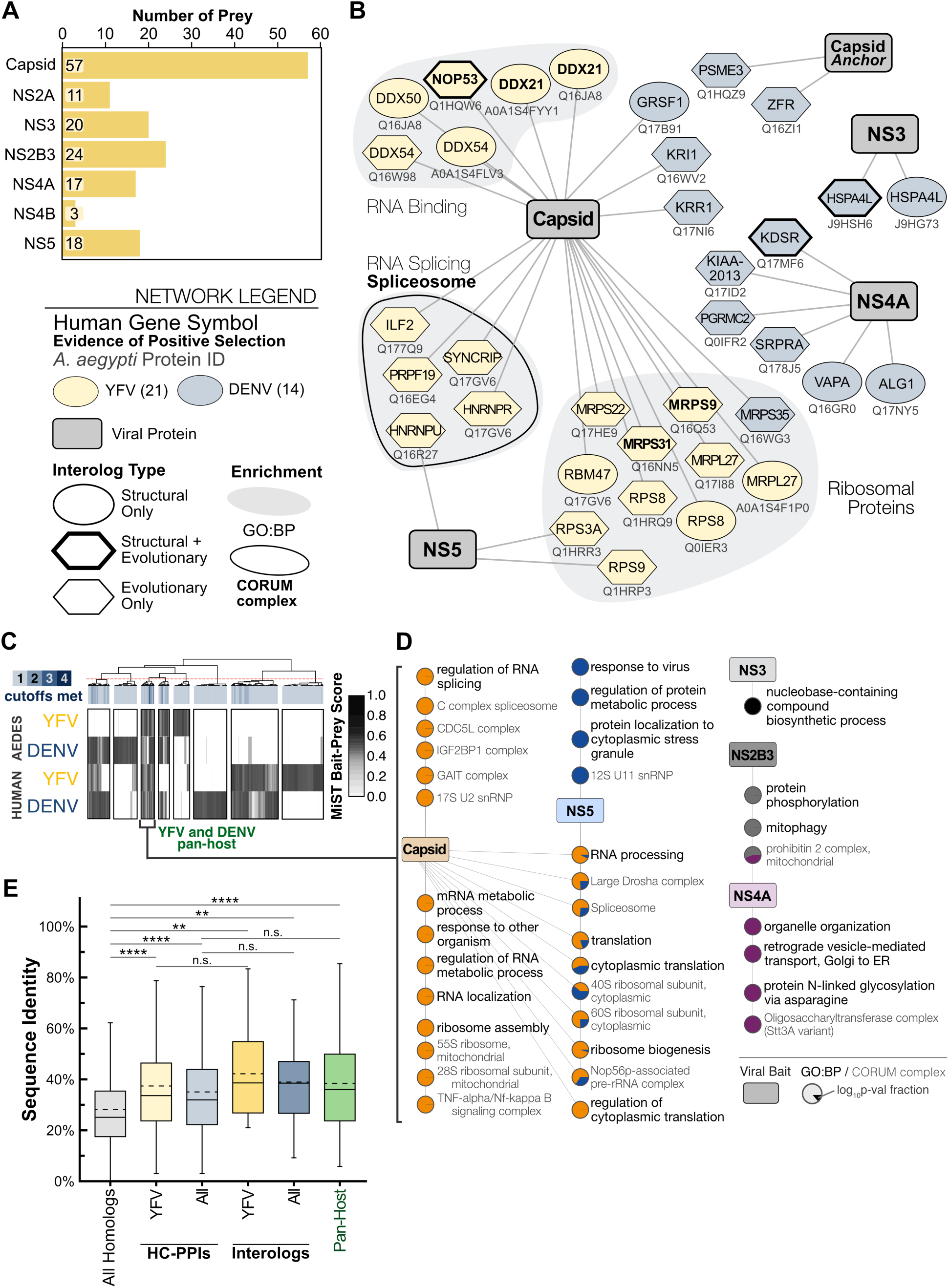
Predicted structures and holistic analysis facilitate interolog identification. (A) Number of HC-PPIs identified per viral bait from mosquito proteomics. (B) Network of structure- and evolutionary-based interologs for both YFV and DENV. The node label is the human gene symbol. The mosquito UniProt ID is listed under the node. Enriched GO:BP categories and CORUM complexes are based on human annotations. Bolded gene IDs indicate proteins found to have positive selection. (C) Hierarchical clustering of flavivirus-mosquito and flavivirus-human PPIs based on MiST scores. Interactions were only considered for viral baits consistently studied across all datasets. An interaction only included if it was a HC-PPI in at least one study and was evaluated for number of studies in which it was considered HC-PPI (cutoffs met shown in blue scale bar). K-means clustering optimized the number of clusters to select a hierarchal cutoff to define clusters (red dashed line). (D) Enrichment network for GO:BP and CORUM complexes from YFV and DENV pan-host cluster, representing interactions conserved across all interactomes analyzed. The color indicates the viral bait associated with the enrichment term and pie sector represents the fractional magnitude of the log_10_p-value of that enrichment. A monochromatic point indicates it was only significant (p < 0.05) for a single viral bait. (E) Box and whisker plots showing the sequence identity values of human-*A. aegypti* homolog pairs of indicated groups of proteins. Solid horizontal lines indicate the median. Dashed horizontal lines indicate the mean. Adjusted p-values were calculated using a Wilcoxon rank-sum test with Bonferroni correction: adjusted p < 0.0001 (****), adjusted p < 0.001 (***), adjusted p < 0.01 (**), adjusted p < 0.05 (*), adjusted p > 0.05 (n.s.).

In addition to host-specific targeting, our previous study with DENV showed that flaviviruses can also target homologous proteins and processes across human and mosquito cells^22^. Our YFV-host proteomics in human and mosquito cells can similarly reveal such interacting homologs (interologs) or interologous processes, utilizing evolutionary relationships between hosts. Comparing YFV-human and YFV-mosquito interactomes revealed 14 interologs at the protein level and an additional 12 complex and pathway-level interologs (Figure S4A).

Given our limited ability to identify direct interologs using evolutionary relationships, we hypothesized that conserved host protein structures might provide a complementary way to measure flavivirus PPI conservation across hosts. To test this, we leveraged recent AI-driven structure prediction and alignment tools to expand interolog identification (Figure S4B). Our workflow used Foldseek^57,58^ and FATCAT^59^ to perform protein structure alignments on the AlphaFold2-predicted structures of the human and *A. aegypti* proteomes. Foldseek and FATCAT output familiar structure similarity scores (*e.g.,* TM-score or RMSD) using complementary approaches. Foldseek uses a 3D alphabet to simplify structural complexity. This enables fast, scalable alignments with large numbers of structures. In contrast, FATCAT flexes proteins into an optimal overlaid structure (superimposition) that maximizes alignment length one residue pair at a time. Combining these analyses enables the simplification of large-scale comparisons into direct, representative alignments without compromising on sensitivity or computational cost.

We first performed proteome-wide rigid structural alignments using Foldseek. To account for potential biases in AlphaFold2 predictions, we supplemented these structural homologs with evolutionary homologs from inParanoid8^60^. Each selected pair then underwent flexible alignment with FATCAT. After overlaying these with the HC-PPIs from YFV and DENV, our structure-based approach increased the number of YFV interologs from 14 to 21 and the number of DENV interologs from 10 to 14 (Figure 4C, Table S5).

However, we were surprised to find no overlap between YFV and DENV interologs. This lack of overlap could be due to fundamentally different dual-host strategies between YFV and DENV. Alternatively, the lack of overlap could stem from technical differences in protein identification and scoring. Indeed, many functionally relevant host interactions may fall below conventional scoring thresholds as “near misses” in one HC-PPI network or another, leading to an underestimation of conserved interactions, processes, and pathways^9,61–64^. To overcome these limitations, we adopted a computational pipeline developed by Gordon and colleagues to evaluate virus-host interactomes holistically beyond strict scoring cutoffs^64^. Across the YFV-human, YFV-mosquito, DENV-human, and DENV-mosquito datasets, we performed hierarchical clustering on MiST scores. We focused our analysis on viral baits studied across all four datasets (Capsid, NS3, NS2B3, NS4A, NS5) and only included data if the bait-prey pair was a HC-PPI in at least one of the four datasets analyzed. With this approach, we found 8 interaction clusters, one of which represents interactions shared across both viruses and both hosts (Figure 4C). This suggests that YFV and DENV indeed share at least some conserved dual-host strategies. Extracting this pan-flavivirus and interologous cluster for enrichment analysis (Figure 4D), we found that individual flavivirus proteins targeted distinct host processes. The Capsid protein targeted translation, ribosome biology, and RNA splicing. NS5 targeted the response to virus and protein localization to the cytoplasmic stress granule, both of which are implicated in flavivirus replication^32,65–69^. NS2B3 interactions were enriched for protein phosphorylation and mitophagy^70,71^. Finally, NS4A expectedly targeted organelle organization, transport, and glycosylation-related pathways. Thus, holistic analysis improves the identification of hijacked pathways fundamental to the dual-host nature of mosquito-borne flaviviruses.

Viruses often exploit these highly conserved host functions to ensure replication, thereby avoiding evolutionary barriers, as hosts also rely on these critical pathways for cellular function. We speculated that targeting these highly conserved systems is especially important for dual-host viruses, such as mosquito-borne flaviviruses, and thus they are targeted as interologs. We compared HC-PPIs and interologs against the entire set of human-mosquito homologous pairs to assess sequence conservation. HC-PPIs showed significantly higher sequence identity compared to homologs in general (Figure 4E, Table S5). This suggests that YFV and DENV evolved to target conserved host proteins, as expected^72^. Our complete set of interologs also showed significantly higher sequence identity compared to homologs in general, yet interologs did not have higher sequence identity than HC-PPIs (Figure 4E, Table S5).

While viruses often target conserved host proteins, as we observed with our HC-PPIs and interologs, these protein interaction surfaces can also contain rapidly evolving sites that serve to break and re-establish virus-host interactions^51–54,56^. So, we explored whether any of the YFV interologs exhibit evidence of participation in this evolutionary arms race. Since there are too few *Aedes* genome assemblies available for analysis, we used simian primate genomes to search for signatures of positive selection (Table S3). *MRPS9* showed strong evidence for positive selection. *DDX21*, *NOP53,* and *MRPS31* showed weaker and less consistent signatures of rapid evolution. Our findings suggest that, although they are enriched for conservation, interologs are not uniquely defined by it, and some may still participate in evolutionary arms races. This highlights the balance between functional constraint and adaptive flexibility that shapes flavivirus-host interactions across host species.

### High-Dimensional Analysis Refines Pan-flavivirus and Divergent Interactions

Previous phylogenetic analysis has shown that YFV is evolutionarily distant from other mosquito-borne flaviviruses, such as DENV, WNV, and ZIKV^20,21^ (Figure 5A). Thus, our YFV dataset provides a powerful foundation for identifying both virus-host interactions specific to YFV and those that are broadly conserved among mosquito-borne flaviviruses. To contextualize these interactions, we compared our YFV-human network to previous studies that used a comparable methodology to identify flavivirus-human PPIs for DENV (serotype 2, strain 16681), ZIKV (strains H/PF/2013 [ZIKVfp] and MR766 [ZIKVug]), and WNV (strain NY 2000-crow 3356)^22,23^. The direct overlap between these datasets only represented ∼5-10% of YFV HC-PPIs (Figure S5A). However, since a holistic analysis that captured “near-misses” was useful for our integration of datasets across multiple hosts (Figure 4C-D), we applied the same technique here. Across the five flavivirus-human PPI datasets, we performed hierarchical clustering on MiST scores for viral baits studied across all four datasets (Capsid, NS3, NS2B3, NS4A, NS5). Again, we performed analysis on bait-prey pairs that were a HC-PPI in at least one of the five datasets analyzed. Our holistic analysis identified 17 interaction clusters (Figure 5B), including conserved interactions across all flaviviruses (cluster 3), YFV-specific interactions (cluster 1), or interactions conserved across all viruses except YFV (YFV-excluded, cluster 8).

**Figure 5.**
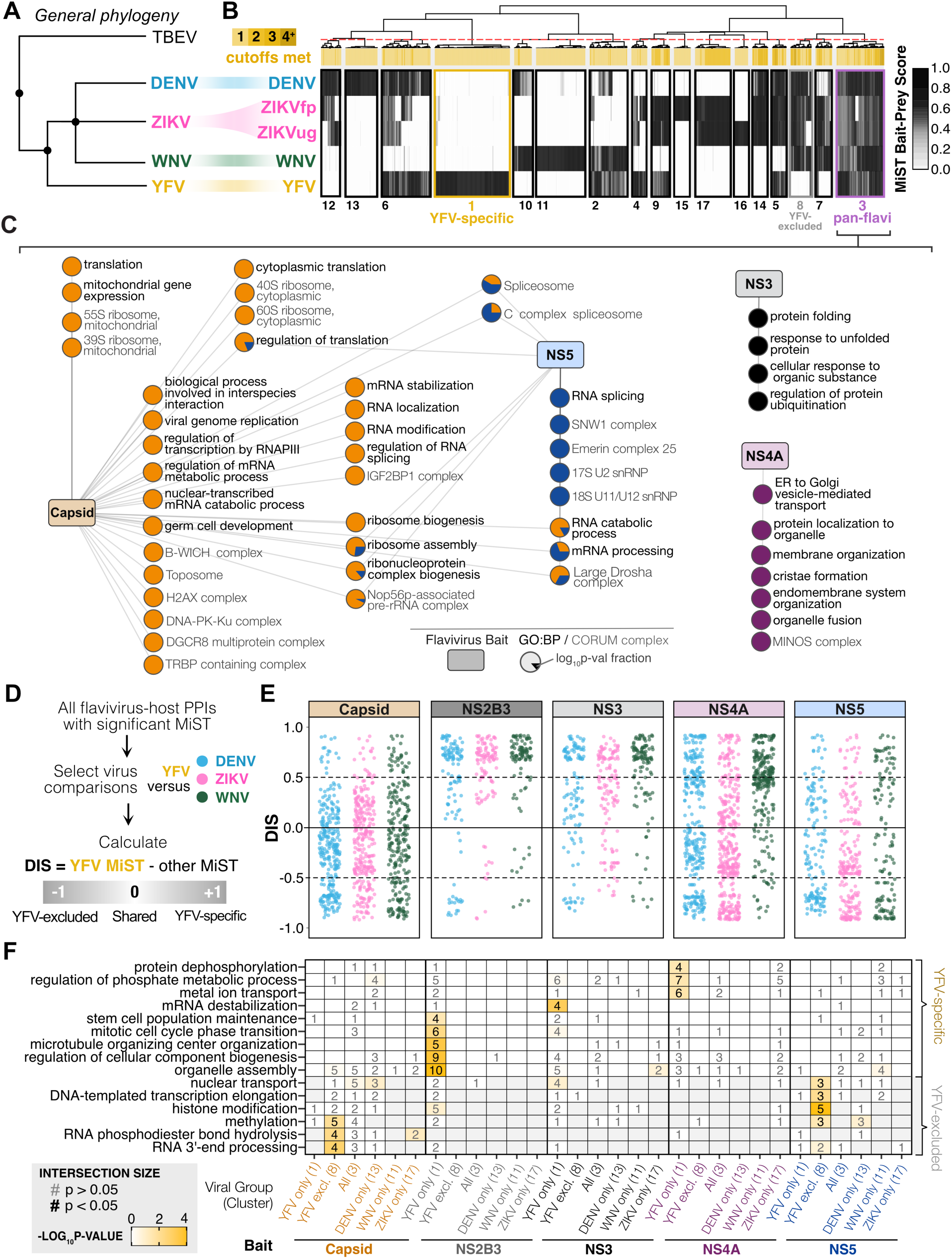
Holistic integration of flavivirus-host interactomes reveals pan-flavivirus and divergent mechanisms of host targeting. (A) Approximate evolutionary relationship between DENV, ZIKV, WNV, and YFV. TBEV was included to represent a true outgroup compared to mosquito-borne flaviviruses. (B) Hierarchical clustering of flavivirus-human PPIs based on MiST scores. Interactions were only considered for viral baits consistently studied across all viruses. An interaction was only included if it was a HC-PPI in at least one study and was evaluated for the number of studies in which it was considered HC-PPI (cutoffs met shown in yellow scale bar). K-means clustering optimized the number of clusters to select a hierarchal cutoff to define clusters (red dashed line). (C) Enrichment network for GO:BP and CORUM complexes from cluster 3, representing pan-flavivirus interactions conserved across all interactomes analyzed. The color indicates the viral bait associated with the enrichment term, and the pie sector represents the fractional magnitude of the log_10_p-value of that enrichment. A monochromatic point indicates it was only significant (p < 0.05) for a single viral bait. (D) Schematic of differential interaction score (DIS) calculation. (E) Plot of YFV DIS distributions, separated by the viral bait. Dot colors indicate the virus used for DIS comparison. (F) Heatmap showing the degree of enrichment (log_10_p-value, yellow scale bar) for unique terms in clusters associated with YFV specifically (cluster 1) or excluding YFV (cluster 8), grouped by viral bait. Enrichment terms were excluded if they showed significant enrichment for other clusters. Other clusters are included in the heatmap to demonstrate the enrichment is unique to clusters 1 or 8. In each tile, the number of interactions associated with that enrichment term is bolded if it was significant (p < 0.05).

To better understand the cellular processes involved in pan-flavivirus PPIs, we performed enrichment analyses on cluster 3 (Figure 5C). Capsid interactions with host proteins dominated this cluster, revealing conserved enrichment of translational processes, mRNA modification, and ribosome biogenesis. Host protein interactions with non-structural proteins were also represented in the conserved interactions. NS3 interactors enriched for protein folding, ubiquitination, and stress response. NS4A interactors enriched for proteins involved in membrane organization and protein transport, reflecting its known role in rearranging ER membranes to form replication complexes^73,74^. NS5 interactors enriched for processes related to RNA splicing, reflecting its known role in perturbing host gene expression^33^. Only NS2B3 interactions did not show enrichment for any biological processes. As with Gordon et al^64^, our analyses confirm that a high-dimensional approach surpasses the limitations of scoring thresholds or single-virus datasets, revealing host targets conserved across these mosquito-borne flaviviruses and related to fundamental aspects of flavivirus biology.

The YFV-specific interactions (cluster 1) and YFV-excluded interactions (cluster 8) suggest an evolution of distinct host hijacking strategies across the flavivirus phylogeny. We therefore evaluated whether these opposing sets of PPIs could be used to understand the evolutionary divergence of YFV relative to other mosquito-borne flaviviruses. To that end, we calculated differences between YFV MiST scores and those of every other flavivirus in the dataset (DENV, ZIKV [we chose to focus on ZIKVfp for simplicity], and WNV). This generated a distribution of differential interaction scores (DIS) relative to YFV for each viral bait. In this YFV-centric approach, a positive DIS indicates the PPI scored highly for YFV relative to the other flavivirus, a negative DIS indicates the PPI scored poorly for YFV, while a near-zero DIS indicates the PPI scored equally in both datasets (Figure 5D).

When visualizing DIS distributions, several trends related to specific viral proteins became apparent (Figure 5E). Capsid interactions were predominantly associated with neutral or negative DIS, suggesting YFV Capsid has many interactions that are shared with other flaviviruses, few YFV-specific interactions, and lacks many of the host interactions observed for other flaviviruses. In contrast, NS2B3 and NS3 exhibited predominantly positive DIS, indicating these proteins have mostly YFV-specific interactions with few shared interactions with other flaviviruses. NS4A and NS5 showed greater variability in their DIS distributions with shared, YFV-specific, and YFV-excluded interactions. We computed Pearson’s correlation coefficients between each pairwise distribution of DIS values and observed consistently high correlations (>0.7, Figure S5B). This suggests YFV’s differential interactions represent *bona fide* divergence in host targeting patterns rather than stochastic deviations in MS detection of proteins in one dataset or another.

To better understand the processes involved in PPIs reflecting YFV’s evolutionary divergence, we extracted statistically significant gene enrichments (p < 0.05) either for the YFV-specific cluster (cluster 1) or the YFV-excluded cluster (cluster 8) (Figure 5E). We excluded any processes that were significantly enriched for other clusters to ensure these processes were truly unique to YFV-specific or YFV-excluded PPIs. These analyses revealed that multiple YFV proteins target host processes in evolutionarily distinct ways. For example, RNA processing and histone modification were significantly enriched for YFV-excluded Capsid and NS5 interactions.

This suggests that although other flaviviruses target these epigenetic processes through Capsid and NS5, YFV does not. Conversely, YFV NS2B3 uniquely targeted host proteins in organelle assembly and microtubule organizing center (MTOC) organization, YFV NS3 uniquely targeted mRNA stability, and YFV NS4A uniquely targeted metal ion transport and protein dephosphorylation.

We extended this analysis to calculate DIS relative to other flaviviruses. This revealed more variable correlations and weaker enrichments for DENV-, ZIKV-, and WNV-specific interactions (Figure S5C-F). This weaker signal is consistent with them being more closely related to each other than YFV and thus having fewer virus-specific or virus-excluded interactions to drive strong DIS scores or enrichment. Together, these results underscore that the pan-flavivirus, YFV-specific, and YFV-excluded clusters represent molecular strategies of host targeting that coincide with the evolution of flaviviruses. To visualize pan-flavivirus and YFV-specific HC-PPIs, we incorporated annotations into our YFV-human HC-PPI visualization using annotations from clusters 3 and 1, respectively (Figure S6).

### Flavivirus Protein Sequence Conservation Does Not Equate to Conservation of PPIs

Protein sequence conservation is often viewed as a predictor of functional conservation^64,75,76^. Structural proteins, such as Capsid, typically exhibit lower sequence conservation than enzymes, like NS5, leading to predictions that structural proteins will retain fewer interactions across viruses. Yet, our preceding analysis using holistic integration and DIS showed a surprisingly high level of conservation for Capsid interactions compared to non-structural proteins (Figures 5C, 5E, and S6). We therefore tested whether the conservation of viral protein sequence can reliably predict flavivirus-host PPI conservation using the four flavivirus datasets (Figure 6A).

**Figure 6.**
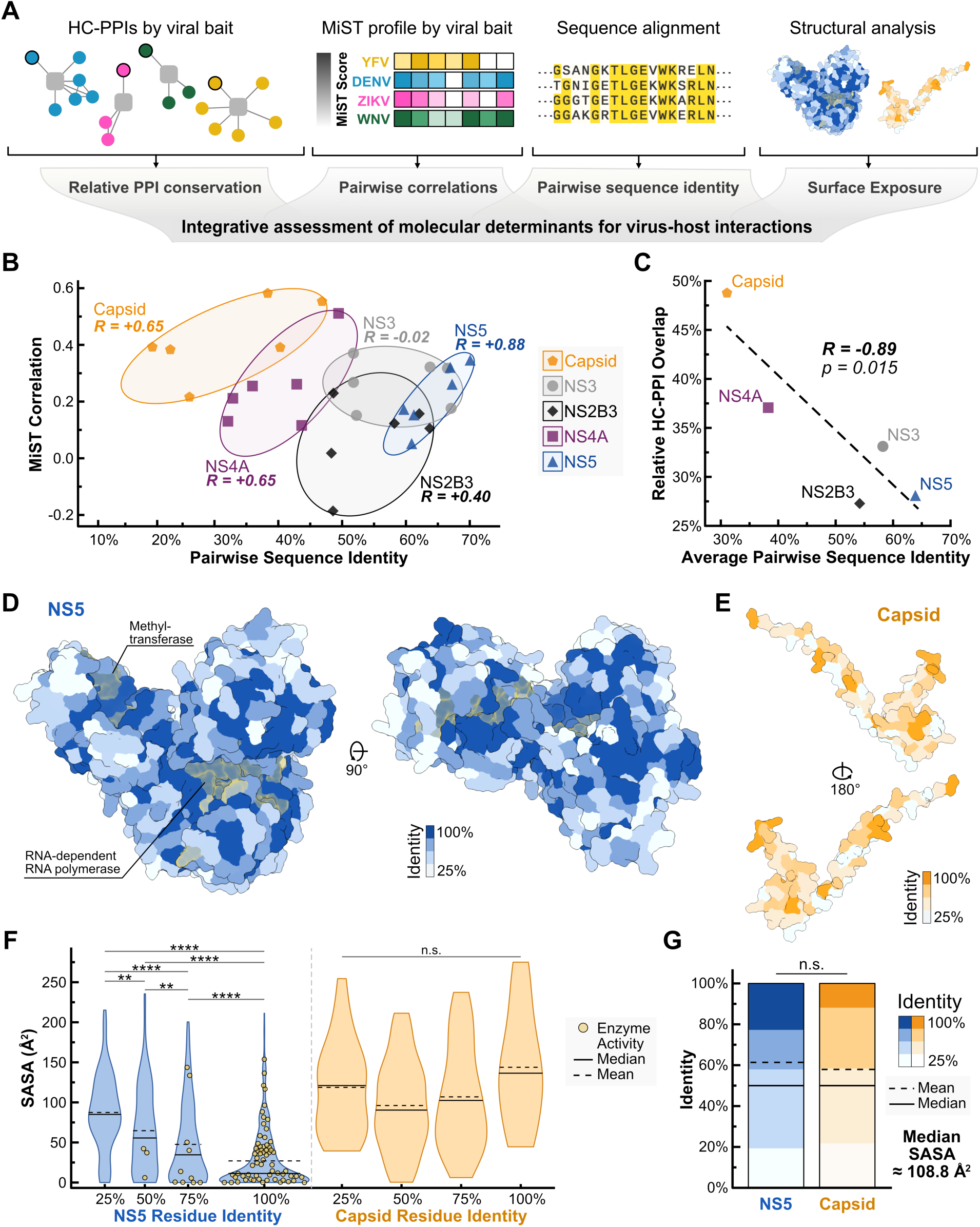
Integration of virus sequence and structure reveals a complex relationship with PPI conservation. (A) Schematic of PPI, sequence, and structural analysis. PPIs were analyzed according to viral baits across all four flavivirus-host PPI datasets. (B) Pearson’s correlation between MiST score profiles and viral protein sequence conservation. For each viral bait, six pairwise correlations were calculated for MiST profiles for each possible pairwise combination between four viruses. The same was done for pairwise sequence identity. Pearson’s correlations between MiST correlations and sequence identity were then calculated for each viral bait and the entire dataset. (C) Pearson’s correlation between relative HC-PPI conservation and viral protein sequence conservation. For each viral bait, mean values of relative HC-PPI conservation and pairwise sequence identity were plotted. (D-E) Visualization of YFV NS5 and Capsid protein structures with protein sequence conservation. Residues with known enzymatic activity are colored with a yellow overlay. Structures were generated using the YFV-Asibi protein sequence and AlphaFold2. (G) Violin plot of solvent accessible surface area (SASA) binned by amino acid conservation for NS5 and Capsid. Yellow dots indicate residues with known enzyme activity. Solid horizontal lines indicate the median. Dashed horizontal lines indicate the mean. Adjusted p-values were calculated using a Wilcoxon rank-sum test with Bonferroni correction: adjusted p < 0.0001 (****), adjusted p < 0.001 (***), adjusted p < 0.01 (**), adjusted p < 0.05 (*), adjusted p > 0.05 (n.s). (H) Stacked bar plot of the distribution of amino acid identity. Only NS5 amino acids with similar SASA as Capsid were considered. P-value was calculated using a Wilcoxon rank-sum (Mann-Whitney U) test: p > 0.05 (n.s).

We calculated pairwise sequence identities and pairwise correlations of MiST scores for each viral bait. Within a single viral bait, we observed a strong positive correlation between these two quantities (Figure 6B). The only exception was NS3. This suggests that the conservation of viral protein sequences predicts conserved virus-host PPIs for individual viral proteins. However, when comparing across viral baits, we observed a slight but unexpected negative correlation between MiST correlation and sequence identity (R = −0.19). This negative correlation across viral baits was more pronounced when our analysis focused on HC-PPIs (R = −0.89, Figure 6C). Thus, surprisingly, we find that more rapidly evolving proteins exhibit higher PPI conservation relative to viral proteins that evolve more slowly.

We investigated whether specific elements of protein structure and function could explain the inverse correlation between viral protein conservation and their host PPI conservation. We focused on NS5 and Capsid because they represent the extremes of PPI and sequence conservation (Figure 6B-C). We first mapped conservation and enzyme activity onto their structures (Figure 6D-E). NS5 is a large protein with a highly conserved core and enzyme active sites. Conserved amino acids are significantly more likely to be buried in regions of low solvent-accessible surface area (SASA), while variable amino acids have significantly higher SASA and opportunity to engage in PPIs (Figure 6F). In contrast, Capsid is a smaller structural protein with lower overall conservation, and its amino acid variation is evenly distributed across low and high SASA regions (Figure 6F). We hypothesized that the enzymatic core of NS5, with high conservation, low SASA, and limited opportunity to interact with host proteins, could be skewing our comparison of NS5 to Capsid. To provide a fairer comparison, we considered only the 233 residues with the highest SASA from NS5 that provide a median of ∼108.7 Å^2^, almost identical to Capsid’s median SASA of 108.8 Å^2^ (Figure 6G). This group of NS5 residues, representing the most solvent-accessible regions of the protein, closely resembles Capsid in both the distribution of residue identities and mean residue identities: 61% and 58% for NS5 and Capsid, respectively. Despite these similar mean identities near the proteins’ surfaces, NS5 still maintains a lower PPI conservation compared to Capsid. Thus, NS5-host PPIs are highly sensitive to small changes in amino acid identity. Our analysis highlights the power of integrating multiple related virus-host PPI networks using a sequence and structure-informed approach, especially when amending a long-standing hypothesis regarding the drivers of conserved virus-host PPIs.

## Discussion

In this study, we systematically mapped YFV-host interactomes in both human and mosquito cells using AP-MS, thereby expanding on previous work on flavivirus-host interactions. We use our YFV-human interactome analysis (Figures 1-2) and a functional screen (Figure 3) to identify RBBP6 as a novel YFV restriction factor. Seeking to move beyond siloed virus-host networks, we applied orthogonal approaches to analyze the data more holistically and from a structural perspective. We integrated evolutionary sequence-structure relationships (Figure 4) and holistic network integration (Figures 5-6) to identify novel YFV biology, and to relate virus and host evolution to the evolution of virus-host PPIs, thereby providing a comprehensive resource on mosquito-borne flavivirus-host interactions.

RBBP6’s antiviral activity was also recently documented for Ebola virus (EBOV)^77^. Our identification of RBBP6 as a YFV restriction factor suggests that this is a restriction factor with panviral potential. Mechanistically, RBBP6 inhibits EBOV by interacting with viral protein 30 (VP30) to displace nucleoprotein (NP) and suppress viral transcription. Similarly, RBBP6 interacts with YFV NS5 and reduces viral genome replication (Figure 3). The RNA-stabilized interaction mechanism is reminiscent of other virus-host molecular interfaces that are also RNA-mediated, such as the interaction between human APOBEC3G and the Vif protein of HIV-1, where RNA might help serve as a ‘molecular glue’^78,79^. The interaction mechanism between NS5 and RBBP6 is distinct from its interaction with EBOV VP30, which binds RBBP6 via a short PxY motif in the Pro-RS domain of RBBP6^77^. However, RBBP6 may still serve as a competitive binder to NS5 to displace NS3, as it does with NP binding to VP30.

Our YFV interactome also provides a starting point for dissecting molecular mechanisms of viral pathogenesis. For example, RBBP6 pathogenic variants increase the risk of myoproliferative neoplasms, which can involve hemorrhaging, thrombocytosis, and coagulopathy^80,81^, similar to severe YF. This link between RBBP6 and hereditary and viral disease is reminiscent of previous findings connecting hereditary and ZIKV-induced microcephaly^22,82^. Extending this beyond a single interaction, an integrative framework enables broader comparisons not only across related viruses but also across shared pathologies. YFV and EBOV share viscerotropic and hemorrhagic pathologies^83^ that may be derived from shared interactions with host proteins, such as RBBP6. Indeed, when comparing HC-PPIs from YFV and EBOV datasets, we identified 21 shared interactions—a statistically significant overlap (p = 8.5e-17, Fisher’s exact test, Figure S7). These findings highlight how viruses with overlapping tropism and pathogenesis may be driven by co-opting similar host factors, even when the viruses are highly evolutionarily divergent from each other.

On the host side, a network-based holistic clustering of YFV and DENV datasets identified interologs conserved across both viruses (Figure 4). This overlap was not revealed through traditional analysis of HC-PPIs, and reinforces the advantages of a holistic analysis that is not subject to strict scoring thresholds. The positive selection signature of interologs, especially mitochondrial ribosomal proteins, is also intriguing given their interaction with flavivirus Capsid in multiple datasets, enrichment in our interolog analysis (Figure S4D), and established roles in promoting the replication of other viruses^84^. Thus, the positively selected genes *MRPS9* and *MRPS31* may be involved in dual-host hijacking, resulting in an unanticipated evolutionary conflict between flaviviruses and their hosts.

We extended our holistic network integration to identify pan-flavivirus and YFV-specific interactions through high-dimensional network integration and differential scoring. We quantified how viral proteins exhibit distinct evolutionary trajectories, retaining some interactions across viruses while diverging in others. This approach was first developed and applied to coronaviruses and different strains of influenza A virus (IAV), revealing the convergence and divergence of viral pathogenic mechanisms^62,64,76^. Our application of this method highlights how binary interaction mapping can benefit from multidimensional data analysis and viewing virus-host protein interactions through the lens of virus evolution.

Finally, by integrating viral protein and PPI conservation with viral protein structure, we revealed distinct insights into flavivirus-host interactions and their potential for evolution. A long-standing assumption in molecular virology is that conserved viral proteins inherently maintain conserved host interactions. Indeed, recent work on other virus families reinforced the expected positive correlation between the conservation of protein sequence and interaction across distinct viral proteins^64,72,76^. Our findings challenge this long-standing assumption, exemplified by the contrast between NS5 and Capsid evolutionary and PPI conservation (Figure 6). Such a divergence suggests that although NS5 is slowly evolving, it can nevertheless rapidly acquire new interactions since small changes in surface-exposed amino acids of NS5 lead to large changes in its protein interactions. Conversely, despite its rapid changes, Capsid is still able to retain conserved PPIs. Whether flaviviruses are unique in this regard remains an open question that can only be answered by continued comprehensive studies of virus-host PPIs.

In summary, we elucidated specific mechanisms of YFV replication as well as the impact of both flavivirus and host evolution on flavivirus-host protein interactions, all stemming from two core proteomics datasets. Yet, this was only possible through the integration of multiple comprehensive flavivirus-host PPI studies and the application of complementary analyses to combine functional assays, virus and host evolution, protein structure prediction, and holistic network integration. Our work adds to an existing body of literature where proteomics provides a powerful foundation to rapidly unravel how viruses engage with host systems through PPIs^22,24,64,85–89^. Its future potential increasingly lies in its integration with orthogonal approaches^9^. Mosquito-borne flaviviruses exemplify the value of this approach because they must navigate interactions across humans and mosquitoes. In the future, complementary methodologies will therefore be paramount to fully describe such adaptation strategies.

### Limitations of this study

Although our study provides a comprehensive YFV-host interactome with several orthogonal approaches, the following limitations should be considered. First, despite integrating structural modeling to address annotation gaps, interolog prediction remains constrained by biases in available host protein structures and their prediction. This may lead to an under-or over-representation of certain interologs. Second, our functional validation was limited to a subset of high-confidence interactions focusing on YFV replication phenotypes, meaning that biologically relevant PPIs may have been overlooked. This could be extended by holistically analyzing our dataset against other viruses that share broad mechanisms of replication or pathogenesis, including other mosquito-borne flaviviruses or EBOV. Third, our proteomic analysis captures steady-state interactions through the expression of individual viral proteins but does not account for the temporal dynamics of virus-host engagement, including activation of innate immune pathways, which may shift over the course of infection. Future studies incorporating both spatial and time-resolved interactomics during infection could better resolve transient or stage-specific interactions. Additionally, since most experimental work is currently limited to YFV, validating our integrative approach in other flaviviruses and vector-host systems will be essential for the generalizability of our findings.

## Materials and Methods

### Resource Availability

The mass spectrometry proteomics data have been deposited to the ProteomeXchange Consortium (http://proteomecentral.proteomexchange.org) via the PRIDE partner repository^90^ with the dataset identifiers of PXD062491 for the YFV-Human AP-MS study and PXD062539 for the YFV-*Aedes aegypti* AP-MS study. All code is available on GitHub (https://github.com/shahlab247/YFV-human-aedes, https://github.com/jayoung/pamlWrapper). Raw western blot, gel, and microscopy images are available upon request. The Foldseek protein structure and NEEDLE protein sequence alignment output data are available on GitHub (https://github.com/shahlab247/YFV-human-aedes) for both the forward (Human to *Aedes aegypti*) and reverse (*Aedes aegypti* to Human) searches. Due to file size constraints, FATCAT structural alignment data are available upon request. Numerical data associated with the alignments for all three methods are available in Table S5. Plasmid maps for the codon-optimized constructs are available upon request.

### Experimental models and tools

#### Cells

HEK293T (from Sam Diaz-Munoz), HeLa (from Luc Snyers), stably expressing Huh7-dCas9 and Vero (ATCC) cells were cultured at 37°C with 5% CO_2_ in Dulbecco’s modified Eagle’s medium (DMEM) (Gibco) supplemented with 10% heat-inactivated fetal bovine serum (HI-FBS; Standard, Gibco). Aag2-AF5 cells^55^ (from Kevin Maringer) were cultured at 28°C with no CO_2_ in Liebowitz’s L-15 media (Gibco) supplemented with 10% FBS (Performance Plus, Gibco), 10% tryptose phosphate broth (Sigma Aldrich), 1X Non-essential amino acids, 1% penicillin-streptomycin (Thomas Scientific), and 1% Amphotericin B (Gibco). All cell lines were tested monthly for mycoplasma contamination using a MycoStrip test kit (Invivogen).

#### Plasmid Constructs

Yellow fever virus isolate Asibi (GenBank accession: KF769016.1) protein-coding sequences were codon-optimized (http://atgme.org) based on codon usage tables for *Homo sapiens* and *Aedes aegypti* (Kazusa DNA Research Institute). Gene fragments were synthesized with Kozak sequences (Twist Bioscience) for Gibson Assembly into pcDNA4_TO with a C-terminal 2xStrep II affinity tag for expression in human cells and pAc with a C-terminal 2xStrep II affinity tag for expression in Aag2-AF5 mosquito cells. HA-tagged RBBP6 domain deletions were previously described (from Chris Basler)^77^. YFV NS5 truncations were designed based on known structural elements and codons optimized for expression in human cells. Gene fragments were synthesized (Twist Bioscience) and ligated into pcDNA4_TO cut with EcoRI and XhoI with C-terminal 2xStrep II affinity tags by Gibson Assembly. Protein coding sequences are provided in Table S6.

#### Viruses

YFV-17D (from Dr. Lark Coffey) stocks were propagated in Vero cells and monitored for CPE. The supernatant was then harvested, and cell debris was removed by centrifugation (Eppendorf centrifuge 5810 R, Rotor S-4-104, 211 g, 5 minutes, 4°C). Cleared supernatant was then distributed into 1 mL aliquots and frozen at −80°C. Each aliquot was used only once to prevent repetitive freeze-thaw cycles. Aliquots were titered by plaque assay (method below).

#### Immunofluorescence microscopy

HeLa cells cultured on #1.5 coverslips were fixed with 4% paraformaldehyde (Fisher) for 15 minutes at room temperature. Cells were permeabilized with 0.1% Triton X-100 (Integra) for 10 minutes and blocked with 5% goat serum (Sigma) in PBS-Tween (0.1% Tween-20, Fisher). Coverslips were incubated with primary antibodies overnight at 4°C. Coverslips were then washed in PBS-Tween and incubated in secondary antibody at room temperature for 1 hour. Nuclei were visualized with Hoechst (1:10000, Invitrogen). Confocal images were acquired using a Zeiss Airyscan LSM980 with Axiocam using a 63x/NA1.40 oil immersion lens. Laser lines at 405, 488, and 543nm were employed sequentially for each image using optics and detector stock settings in the “Dye List” portion of the FluoView microscope-controlling software. All antibodies and dilutions are listed in Table S7. Microscopy images were analyzed using ImageJ (Fiji) software^91^. Signal colocalization was quantified using Pearson’s correlation coefficient (R-value) determined with the “JACoP” analysis tool imported into Fiji after setting regions of interest around entire individual cells across at least 5 images.

#### Western blotting

For whole cell lysates, cells were lysed in RIPA buffer (150 mM NaCl, 50 mM Tris Base, 1% Triton X-100, 0.5% sodium deoxycholate) supplemented with protease inhibitors for 5 minutes at room temperature. Otherwise, cells were lysed in IP buffer supplemented with Complete-EDTA-free Protease Inhibitor Cocktail (Sigma Aldrich, USA). Cell lysate was incubated on ice for 30 minutes prior to centrifugation (Eppendorf centrifuge 5424 R, Rotor FA-45-24-11, 13,500 g, 4°C, 20 min). When possible, the total protein concentration of each sample was normalized by BCA assay. Protein samples (lysates or IP eluates) were resuspended in NuPAGE LDS sample buffer supplemented with TCEP and boiled at 95°C for 10 minutes. Samples were run on 7.5-12% polyacrylamide gels for ∼1 hour at 150 V and transferred to PVDF membranes (VWR) for 1 hour at 330 mA on ice. Membranes were then blocked in 5% milk solution for 1 hour prior to overnight incubation in primary antibodies (Table S7) at 4°C. Membranes were washed 3x in Tris-buffered saline with Tween-20 (TBS-T) (150 mM NaCl, 20 mM Tris Base, 0.1% Tween-20, Fisher) and incubated with HRP-conjugated secondary antibodies in 5% milk for 1 hour at room temperature. Membranes were again washed 3x in TBS-T and 1x in TBS (without Tween-20) prior to Pierce^TM^ ECL activation (Fisher). Membranes were imaged using Amersham Imager 600 (GE).

#### Plaque assay

Vero cells were seeded at 0.8×10^6^ cells/well and grown in 6-well dishes overnight. Virus aliquots were thawed on ice and subjected to 6-fold serial dilution. Media was removed from Vero cells, and the monolayer was washed with 1xD-PBS. For YFV-17D, 900 µL of each virus dilution was then added to the cells and incubated for 4 h at 37°C. Virus was then removed, and cells were overlayed with 2 mL of DMEM with 0.8% methylcellulose (Sigma), 1% FBS, 1% penicillin/streptomycin, and incubated at 37°C for 7 days. Cells were then fixed with 4% formaldehyde and stained with 0.23% crystal violet solution (Fisher) for 30 minutes at room temperature. The solution was then removed, and plaques were counted.

### Generation of YFV proteomic networks

#### Expression of viral proteins

For each affinity purification (AP) [YFV-Asibi protein constructs, empty vector, green fluorescent protein (GFP)], 1×10^7^ human HEK293T or 3×10^7^ Aag2-AF5 mosquito cells were plated in 15 cm cell culture dishes 24 hours prior to transfection. A total of 20 μg of plasmid DNA was transfected in a 1 μg: 3 uL Polyjet (SignaGen) ratio for expression in HEK293T cells. Similarly, the same mass of plasmid DNA was transfected in a 1 μg: 2 μL TransIT-Insect (Mirus Bio) ratio for expression in Aag2-AF5 cells. Forty hours post-transfection, cells were gently washed in D-PBS and dissociated with 10 mM EDTA in D-PBS. Cells were pelleted (250×*g*) at 4°C for 5 min, washed in D-PBS, re-pelleted and lysed for 30 minutes on a rotator at 4°C in AP Buffer [50 mM Tris-HCl pH 7.4, 150 mM NaCl, 1 mM EDTA, Pierce EDTA-free protease tablets (Thermo Scientific)] supplemented with 0.5% Nonidet P 40 Substitute (NP40; Fluka Analytical). Subsequently, cell debris was pelleted (4500×*g*) at 4°C for 20 minutes. A fraction of lysate was resolved on a hand-cast 4⟶20% gradient SDS-PAGE gel to evaluate 2xStrep-tagged protein expression by Western blotting. The remaining lysate was used for subsequent affinity purification.

#### Strep-tag affinity purification and digest

Each cell lysate was combined with 100 μL of a 50% Strep-Tactin Sepharose bead (IBA Lifesciences) slurry and incubated on a tube rotator for 2 hours at 4°C. After binding, beads were washed in three times AP Buffer supplemented with 0.05% NP-40. For each wash, 1 mL of AP buffer was added, tubes were inverted, and beads were pelleted (∼500×*g*) at 4°C before removing the supernatant. After a final wash in AP buffer without NP-40, 1/3 of the beads were transferred to a fresh tube and eluted in 25 μL of 2.5 mM D-Desthiobiotin (IBA Lifesciences) by agitation (1300 rpm) at room temperature. Like cell lysates, this eluate was resolved by SDS-PAGE and protein expression was evaluated using both Western blotting and a Pierce Silver Stain (Thermo Scientific). The remaining 2/3 of the beads were equilibrated in diminishing volumes (200 μL, 100 μL, 50 μL, 25 μL) of 50 mM Triethylammonium bicarbonate (TEAB) buffer (Thermo Scientific) at 4°C for 15 minutes on a tube rotator. At the final wash, 500 ng of Sequencing Grade Modified Trypsin (Promega) was added to each sample. Beads were then incubated under agitation (1300 rpm) overnight at 37°C. The following morning, the digest supernatant was transferred to a new protein LoBind tube (Eppendorf). Another 200 ng of trypsin was added to the beads, along with 40 μL of TEAB, and incubated for an additional 2 hours; this digest was then combined with the previous extract.

#### Mass spectrometry

Digested peptides were directly loaded onto an Evosep C18 tip and separated using the Evosep One and the standard (human samples) or 100 spd High Organic method (mosquito samples) (Evosep). Peptides were eluted and ionized using a Bruker Captive Spray emitter. A Bruker TimsTof Pro 2 mass spectrometer running in diaPASEF mode was used for acquisition. The acquisition scheme used for diaPASEF consisted of 6×3 50 m/z windows per PASEF scan. DIA data was searched using Spectronaut 17 (Biognosys) against the human (UP000005640, downloaded 6/15/2022) or *Aedes aegypti* (UP000008820, downloaded 2/10/2023) UniProt proteomes, YFV protein sequences, and the standardized common contaminants database. The Direct DIA workflow was used under default settings. Briefly, trypsin/P Specific was set for the enzyme, allowing two missed cleavages. Fixed Modifications were set for Carbamidomethyl, and variable modifications were set to Acetyl (Protein N-term) and Oxidation. For DIA search identification, PSM and Protein Group FDR were set to 1%. A minimum of 2 peptides per protein group were required for quantification. Raw intensity values were normalized to all mapped peptides. Background detections were eliminated by excluding intensity values below the median intensity value for each protein, and common contaminants were removed.

#### PPI scoring and network visualization

We scored protein interactions with both Mass spectrometry interaction STatistics (MiST)^92^ and Significance Analysis of INTeractome Express (SAINTexpress)^93^ platforms. For MiST scoring, MS1 intensity values were treated as the quantifying feature, and weights were set to R= 0.35, S = 0.57, and A = 0.08, in line with our previous flavivirus work^22^. Samples or baits were removed from further analysis if they exhibited poor clustering or minimal/no detection of the bait itself in the quality control step. Specificity exclusions were applied to baits with shared sequences (prM-prME, NS2B-NS2B3, NS3-NS2B3) or those that exhibited high PPI overlap across previous flavivirus-host proteomic studies (NS4A-NS4B and NS4A-NS2A). Each bait was compared to at least 8 other baits for specificity scoring regardless of these exclusions. Concurrently, protein interactions were also scored with SAINTexpress using the Galaxy APOSTL server (http://apostl.moffitt.org/) under default parameters, with the GFP bait serving as the source of control counts.

We mirrored a two-step filtering strategy for identifying high-confidence interactions previously^64,94^. First, we selected all protein interactions with a MiST score > 0.67—as this threshold was rigorous in past flavivirus-host proteomic studies^22^—and a SAINTexpress score > 0.95 and Bayesian false-discovery rate (BFDR) ≤ 0.05. We then identified if these proteins were part of a curated human CORUM (4.0 release) protein complex^38^, recovering any additional complex members with a MiST score ≥ 0.6, a SAINTexpress score > 0.95 & BFDR ≤ 0.05, and interacting with the same viral bait. All PPI networks were built in Cytoscape (v3.9.0) and formatted with Affinity Designer. Prey proteins were annotated for GO: Biological Processes (GO:BP), protein complexes were extracted from CORUM, and additional or overlapping annotations were manually curated and grouped as needed.

### Phenotypic screening and RBBP6 assays

#### CRISPRi knockdown

Custom synthetic guide RNAs (gRNA) were acquired from Sigma and resuspended to 3 µM in TE buffer (10 mM Tris Base, 1 mM EDTA, pH 8.0). For the knockdown experiment, 375 μL of Opti-MEM (Life Technologies, USA), 5 μL of TransIT-CRISPR transfection reagent (Sigma, USA), and 20 µL of gene-specific sgRNAs were combined and incubated at room temperature for 20 min, then added to the appropriate well of the 12-well plate. Approximately 0.5 × 10^5^ cells were plated to corresponding 12 well plates. After 72h, cells were washed with 1xPBS, lysed in RNA lysis buffer (Thomas Scientific, USA), and the efficiency of gene knockdown was estimated by qRT-PCR. Single-stranded cDNA was synthesized from total RNA using the iScript cDNA Synthesis Kit (BioRad, USA) according to the manufacturer’s recommendations. qPCR was performed using SYBR Master mix (Invitrogen, USA). All qRT-PCR data were normalized to the expression of the *GAPDH* gene. Expression fold changes were calculated by using the ΔΔCt method. Genes with knockdown greater than 50% were prioritized for virus replication assays. gRNA and qPCR primer sequences are available in Table S8.

#### CRISPRi infection assay

Cells were infected with YFV-17D at the indicated MOI 72h after CRISPRi knockdown. Supernatant aliquots were harvested at 0, 24, 48, and 72 hpi and frozen at –80°C. The titration of viral load in the infectious culture supernatants collected at different time points was conducted by plaque assay in Vero cells in triplicates. For the CRISPRi screen, robust *z*-scores were calculated for log-transformed titers measured at 48 hpi. The complete screening results can be found in Table S2.

#### Endogenous RBBP6 immunoprecipitation

At 48 hours post-infection, mock or YFV-17D-infected Huh7 cells (MOI of 1) were collected, washed twice with ice-cold PBS and lysed with immunoprecipitation buffer (50 mM Tris-HCl pH 7.5, 140 mM NaCl, 1.5 mM MgCl_2_, 0.5% NP-40) supplemented with Complete-EDTA-free Protease Inhibitor Cocktail. Cell lysates were incubated for 30 min on ice. After clearing lysates by centrifugation at 1000 x g for 10 min at 4°C, they were treated or not with RNaseA (Thermo Fisher Scientific, USA) for another 30 min. Following this, 25 μL of lysates (treated and not treated with RNaseA) were preserved as protein Input while another aliquot (25 μL) was used to perform IP. A total of 25 µl of Protein A Mag Sepharose beads (Cytiva, USA) had been equilibrated with cold washing buffer (50 mM tris, 150 mM NaCl pH 7.5) and incubated overnight with agitation at 4°C with anti-RBBP6 (Origene, USA) and negative control rabbit IgG (Santa Cruz Biotechnology, USA) antibodies. Following this, the cell lysates were added to the beads and incubated for 1 h on ice. The supernatants were removed, and beads were washed 3 times with 500 µl washing buffer (50 mM tris, 150 mM NaCl pH 7.5). After the last washing step, beads were resuspended in 50 µl of western blot loading buffer and boiled at 95°C for 10 min. Obtained samples (IPs and Inputs) were analyzed by Western blotting.

#### YFV replicon assay

YF-Rluc-2A-WT expressing plasmid (gift of Richard Kuhn^50^) (10 μg) was linearized using XhoI enzyme (NEB, United States) prior to *in vitro* transcription. The linearized DNA was purified using Gel and PCR Clean-up, NucleoSpin (Macherey-Nagel, USA). The capped RNA transcripts were synthesized using an SP6 mMessage mMachine kit (Invitrogen, USA), and the obtained RNA was precipitated with Lithium Chloride following the manufacturer’s instructions. Huh7-dCas cells were first transfected with ncgRNA, RBBP6 gRNA1, and RBBP6 gRNA2 and 48h later transfected with the obtained RNA transcripts (5 μg/per reaction) using Lipofectamine 2000 (Invitrogen, USA). Briefly, transfection mixtures were incubated at room temperature for 15 min, added to the cells, and then incubated for 4–6 h at 37°C. After incubation, 1mL of complete growth medium was added to each well. The cells were lysed at 0, 3, 6, 12, 24, and 48 h post-transfection and stored at −20°C. All the measurements of luciferase activity were performed by *Renilla* luciferase assay kit according to manufacturer instructions (Promega Corporation, USA) and using a Spectramax m2 plate reader (Molecular Devices, USA).

### Identification and conservation analysis of interologs

#### Interolog mapping of mosquito proteins

To appropriately compare human and mosquito PPI networks, we mapped *Aedes aegypti* mosquito proteins to human orthologs. We leveraged a multi-stepped approach to maximize ortholog identification. In brief, we extracted all possible ortholog mappings from the InParanoid 8 database^60^ between *Homo sapiens* and *Aedes aegypti*, mapping a majority of mosquito proteins. Some annotations were additionally mapped using OrthoDB^95^. Each of these annotations is provided in the supplemental network file (Table S4B).

#### Identification of interologs by AlphaFold2 and Foldseek

To enhance interolog identification, we extracted both *A. aegypti* and human proteomes from the AlphaFold EMBL-EBI database (https://alphafold.ebi.ac.uk/). The human proteome is readily available for download, while the *A. aegypti* proteome must be downloaded from a separate repository available through the same website. In total, there were 23,391 and 30,561 structures for each organism, respectively. These numbers exceed the actual quantity of annotated and predicted proteins for each organism because larger proteins are split into sections to make predictions via AlphaFold tractable. We first performed an all-versus-all comparison of the two proteomes using Foldseek to identify potential structural homologs for every protein: human to *A. aegypti* (forward search) and *A. aegypti* to human (reverse search). Foldseek performs a rigid structural alignment with a 3D alphabet approach that allows for rapid searches against entire proteomes on local computer systems. Foldseek can identify hundreds of similar structures for a single query with varying levels of confidence structure (represented as an e-value), so we limited results to any structure with an e-value below 1e-10. We then took the intersection of the two datasets for an aggregate of structural homolog pairs between the forward and reverse search of the human and *A. aegypti* proteomes. From this list, the 3 pairs with the smallest e-values were retained for further analysis. We added homologs identified by inParanoid8 to alleviate bias toward highly structured proteins that could be predicted confidently by AlphaFold. We performed flexible structural alignments using FATCAT for all filtered structural homolog pairs. Duplicate pairs generated from proteins with multiple sectioned structures were removed using the p-value output by FATCAT, resulting in a final 32,749 structural homolog pairs. These pairs were compared to both the YFV and DENV HC-PPI datasets. If either a human or *A. aegypti* protein appeared as a HC-PPI with a viral bait, we classified the pair as a HC-PPI. If both the human and *A. aegypti* proteins in a structural homolog pair appeared as a HC-PPI for the same viral bait, we classified them as structural interologs.

#### Sequence similarity analysis of interologs

For the 32,749 structural homolog pairs, we performed sequence alignments using the EMBOSS Needle algorithm. We maintained the same categorizations for each pair plotted.

### Positive selection analysis

To examine the evolutionary selective pressures acting on *RBBP6* and 17 unique high-confidence interologs in primates, we first selected the human splice isoform with the longest ORF for each gene. We used each ORF as query in a blastn^96^ search of NCBI’s nt database, restricting hits to simian primates. For each non-human primate species, we collected the hit with the highest bit score and obtained its ORF sequence. We generated an in-frame alignment using the MACSE^97^ algorithm and made manual adjustments to fix likely gene prediction errors and alignment ambiguities. We then estimated a phylogeny from each alignment using phyml^98^, with the GTR nucleotide substitution model, four rate categories, and estimating the proportion of invariable sites.

We used the resulting alignments and trees as inputs to the PAML^99^ algorithm (codon model 2, starting omega 0.4), comparing the likelihoods of evolutionary models that allow positive selection at a subset of sites (models 8 and 2) with nested models that allow only sites with purifying or neutral selection (models 7, 8a and 1). Results are shown in Table S3. Four genes had a p-value ≤ 0.05 in our preferred test (model 8 versus 8a); however, after correction for multiple testing using the Benjamini and Hochberg (BH) method^100^, only one (*MRPS9*) remained significant.

We also ran the GARD algorithm^101^ to examine alignments for recombination and gene conversion, which can cause false positives in evolutionary analysis. One of the genes that showed evidence of positive selection (*NOP53*) also showed evidence of recombination. We reran PAML on each GARD-identified segment separately, finding that the signal of positive selection remained for segment 1. However, close inspection of the *NOP53* results revealed only a single amino acid position (codon 6) with a high likelihood of positive selection (> 90% BEB posterior probability) - this codon overlaps a CpG dinucleotide in a very CpG-rich region at the 5’ end of the gene. Methylated CpG sites are known to mutate at much higher rates than other sequences^102^, a phenomenon that PAML (like other algorithms to detect selective pressure) does not account for. While it is possible that positive selection drove changes at this residue, it seems more likely that the number of changes observed at this site reflects its hypermutability. As a conservative approach to mitigate the impact of CpG hypermutation, we masked alignments to remove any codon that overlapped a CpG dinucleotide in any species in the alignment and re-ran PAML. Statistical support for positive selection disappeared for *NOP53* but remained for *MRPS9*, *DDX21*, and *MRPS31*. We also checked whether the signature of positive selection in these four genes was robust to the use of alternative starting parameters (codon models 2 or 3, starting omega 0.4 or 3, model 8 versus 8a p ≤ 0.05). For *MRPS9*, *DDX21*, and *MRPS31*, all four starting parameter combinations yielded evidence of positive selection; *NOP53* gave more equivocal results, where two of the four-parameter combinations had p ≤ 0.022, and the other two had p = 0.059. Full positive selection results are available in Table S3.

### Integrative analysis of protein interactions

#### Network similarity metrics

When comparing the overlapping bait-prey data between various datasets, the relevant data were reformatted into a contingency table. A Fisher’s exact test was performed to assess the statistical significance of the overlap using the GeneOverlap R package (v1.32.0). Venn diagrams showing the degree of overlap were created with the BioVenn R package (v1.1.3).

#### Enrichment analyses

All gene enrichment analyses were conducted with the R package gprofiler2 (0.2.1). Typically, we separated genes into lists by viral bait and/or hierarchal cluster prior to enrichment. Using the “gost” function, we tested for functional enrichment across Gene Ontology (GO), Reactome Pathways, and CORUM databases. Statistical significance was calculated with a hypergeometric test, correcting for false discovery rate (FDR) using the Bonferroni method. Enrichment analyses for both human and mosquito data were performed using *Homo sapiens* databases against all annotated genes, meaning mosquito proteins without an annotated ortholog were excluded from these analyses.

#### Hierarchal clustering of protein interactions

We performed hierarchal clustering on interactions for viral bait proteins shared across all five viral bait datasets and that met their respective MiST cutoffs in at least one of the datasets. Clustering was performed on MiST scores for each virus-host interaction using the “hclust” function and “ward.D2” method of the base R package stats. We optimized the number of clusters by applying k-means clustering and selecting a hierarchal cutoff that produced the equivalent number of clusters. Enrichment analyses were conducted by both cluster and viral protein, as described previously. Thus, we could extract unique or shared enrichment categories (p < 0.05) for a particular bait and cluster. Clustering was visualized with the R package pheatmap (v1.0.12).

#### DIS analysis and scoring

Differential interaction scores (DIS) were calculated for the same subset of data used for hierarchal clustering by finding the difference in MiST scores between two viruses for a particular bait-prey interaction. We excluded cases where both MiST scores were 0. To ascertain the level of correlation across DIS, Pearson’s correlations were calculated based on the distribution of DIS between two particular comparisons. Distributions were visualized with the R package ggplot2 (v3.4.1).

#### Enrichment network visualization

To simplify enrichment data for visualization, we isolated enriched terms between 1 and 2000 genes and eliminated redundancy by collapsing terms with greater than 50% similarity, retaining the term with the lowest adjusted p-value. Cytoscape (v3.9.0)^103^ was used to plot these as networks, and the “geom_tile” function of ggplot2 created heatmaps of relevant baits/clusters.

### Conservation analysis of PPIs by viral bait using protein sequence and structure

#### HC-PPI conservation analysis and MiST profile correlations

MiST scores for YFV, DENV, WNV, and ZIKV were compiled by viral bait, excluding any entries that were not detected by MS in all four datasets. Pairwise Pearson’s correlation coefficients were calculated in R. The union of HC-PPIs for YFV, DENV, WNV, and ZIKV was compiled by viral bait. For each PPI, a relative conservation was calculated based on the fraction of datasets that PPI was conserved. This was summed over the entire union for each viral bait. Since each viral bait had a different number of HC-PPIs, ranging from 20 to 74, this was normalized to the total number of HC-PPIs for that bait.

#### Viral protein sequence identity calculations and visualization

Amino acid sequences for flavivirus proteins from YFV, WNV, DENV, and ZIKV were collected from sequences used in published proteomics studies^22,23^ or this study. Sequence identity was determined using the T-Coffee alignment algorithm within SnapGene (v4.3.11) (www.snapgene.com). Protein structures were predicted using AlphaFold2 and visualized using UCSF ChimeraX (v1.9)^104^. YFV Capsid and NS5 structures were colored based on the four viruses, ranging from 25% (1 out of 4) to 100% (4 out of 4) amino acid identity. Solvent-accessible surface area (SASA) calculations were performed using UCSF ChimeraX (v1.9).

### Statistical analysis

Simple linear regression was applied to data when indicated to calculate Pearson’s correlation coefficients. P-values of correlation coefficients were calculated by bootstrapping the linear regression. The indicated data was shuffled, and the correlation coefficient was calculated. This was repeated 100,000 times to create a distribution of correlation coefficients. The percentage of randomly generated correlation coefficients greater (for positive correlation coefficients) or less (for negative correlation coefficients) than the true correlation coefficient was taken as the P-value.

## Acknowledgements

We thank members of the Shah Lab for constructive feedback and rewarding discussions. Funding for this work was provided by NIH R21AI168716, NIH R01AI170857, and the Laboratory Directed Research and Development Program at Sandia National Laboratories under Project 233120 to P.S.S. Funding to the proteomics core was provided by Dr. Neil Hunter and the Howard Hughes Medical Institute. Work in the Malik lab was supported by NIH U54AI170792 (PI: Nevan Krogan); H.S.M. is an Investigator at the Howard Hughes Medical Institute. The Zeiss 980 confocal microscope used in this study was purchased using NIH Shared Instrumentation Grant S10OD026702. We thank Thomas Wilkop of the MCB Light Microscopy Imaging Facility, which is a UC Davis Campus Core Research Facility, for assistance using this microscope.

## Author Contributions

P.S.S. and M.W.K. conceived the project and designed experiments, with input from L.C., C. L. S. S., H.S.M., J.M.Y., and T.B.. M.W.K. performed affinity purification and designed computational pipeline for scoring YFV proteomics and subsequent comparative analyses with contributions from C.L.S.S. and P.S.S.. L.C. and C.J.F. performed CRISPRi knockdowns and associated virus assays. L.C. performed assays related to RBBP6. M.W.K., L.C., A.T.F., V.P., and C.J.F. performed Western blotting analyses. A.T.F. and M.W.K. conducted molecular cloning, performed immunostaining and microscopy. C.L.S.S., T.B., J.M.Y., and P.S.S. performed computational structure and evolution studies. M.W.K., C.L.S.S, and P.S.S. performed data visualization. P.S.S. and H.S.M. supervised the research. P.S.S. and H.S.M. acquired funding. M.W.K., H.S.M, and P.S.S. wrote The Manuscript. L.C., C.L.S.S., A.T.F., T.B., and J.M.Y. edited The Manuscript.

## Supplementary Files

**Table S7:** Antibodies.

| Antibody | Host Species | Dilution Used | Supplier (Catalog #) | RRID |
| --- | --- | --- | --- | --- |
| GAPDH | Mouse | 1:1000 (WB) | Fisher (PIMA515738) | AB_2537652 |
| FLAG-M2 | Mouse | 1:200 (IF),<br>1:1000 (WB) | MilliporeSigma (F1804) | AB_262044 |
| Strep | Mouse | 1:1000 (WB),<br>1:500 (IF) | Qiagen (34850) | AB_2810987 |
| HA | Mouse | 1:1000 (WB),<br>1:200 (IF) | MilliporeSigma (H3663) | AB_262051 |
| TOMM20 | Rabbit | 1:250 (IF) | ThomasScientific (11802-1-AP) | AB_2207530 |
| Calnexin | Rabbit | 1:250 (IF) | Proteintech (10427-2-AP) | AB_2069033 |
| RBBP6 | Rabbit | 1:1000 (WB) | ThermoFisher (PA5-23054) | AB_11152136 |
| YFV NS5 | Rabbit | 1:1000 (WB) | GeneTex (GTX134141) | AB_2887231 |
| GRASP65 | Rabbit | 1:250 (IF) | From Dr. Nevan Krogan | N/A |
| Anti-Mouse IgG-HRP | Rabbit | 1:5000 (WB) | SouthernBiotech (6170-05) | AB_2796243 |
| Anti-Rabbit IgG-HRP | Goat | 1:5000 (WB) | SouthernBiotech (4030-05) | AB_2687483 |
| Anti-Mouse AlexaFlour-488 | Goat | 1:1000 (IF) | Invitrogen (A28175) | AB_2536161 |
| Anti-Mouse AlexaFlour-555 | Goat | 1:1000 (IF) | Invitrogen (A21422) | AB_2535844 |
| Anti-Rabbit AlexaFlour-488 | Goat | 1:1000 (IF) | Invitrogen (A11008) | AB_143165 |
| Anti-Rabbit AlexaFlour-555 | Goat | 1:1000 (IF) | Invitrogen (A27039) | AB_2536100 |

**Table S8:**
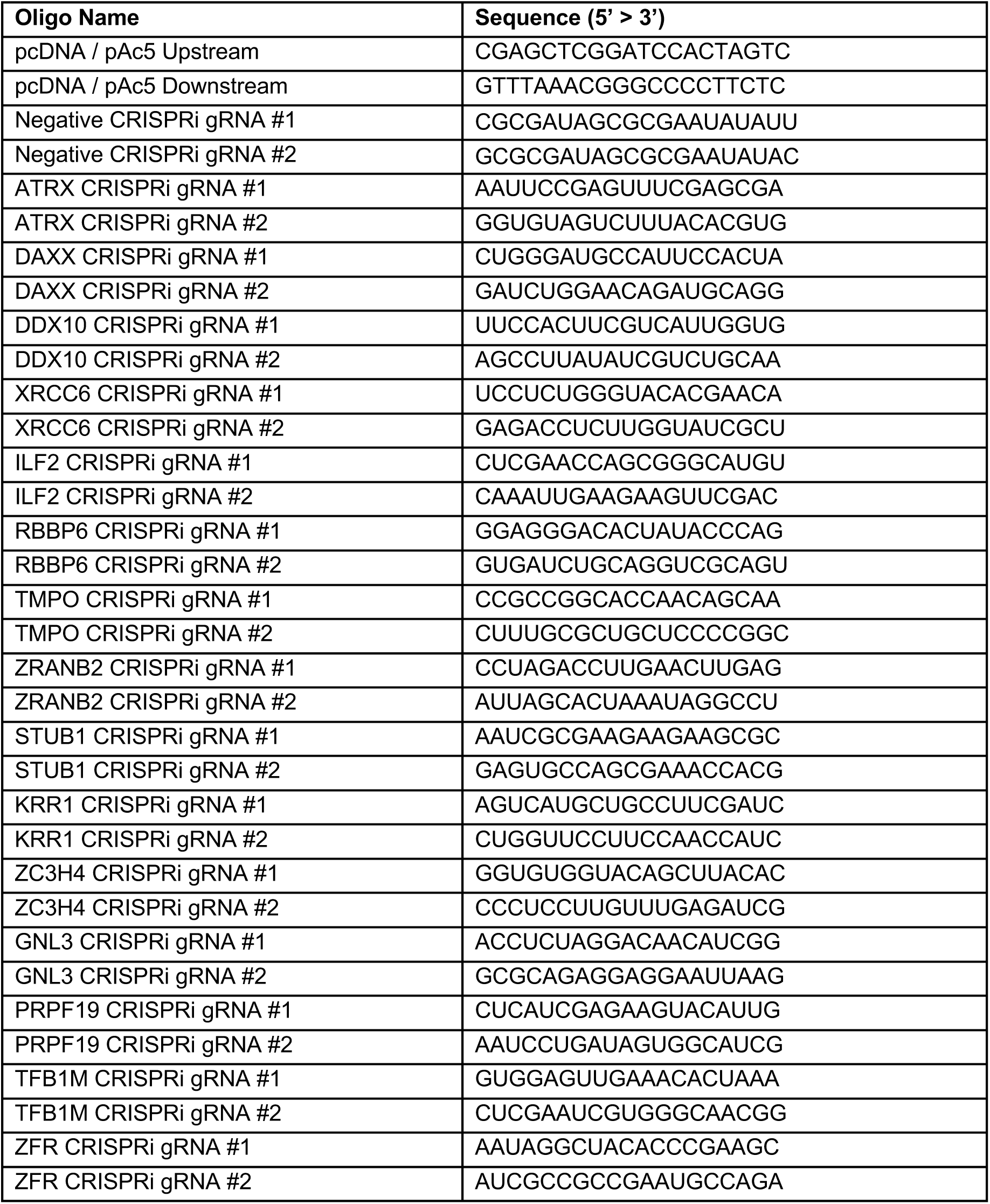
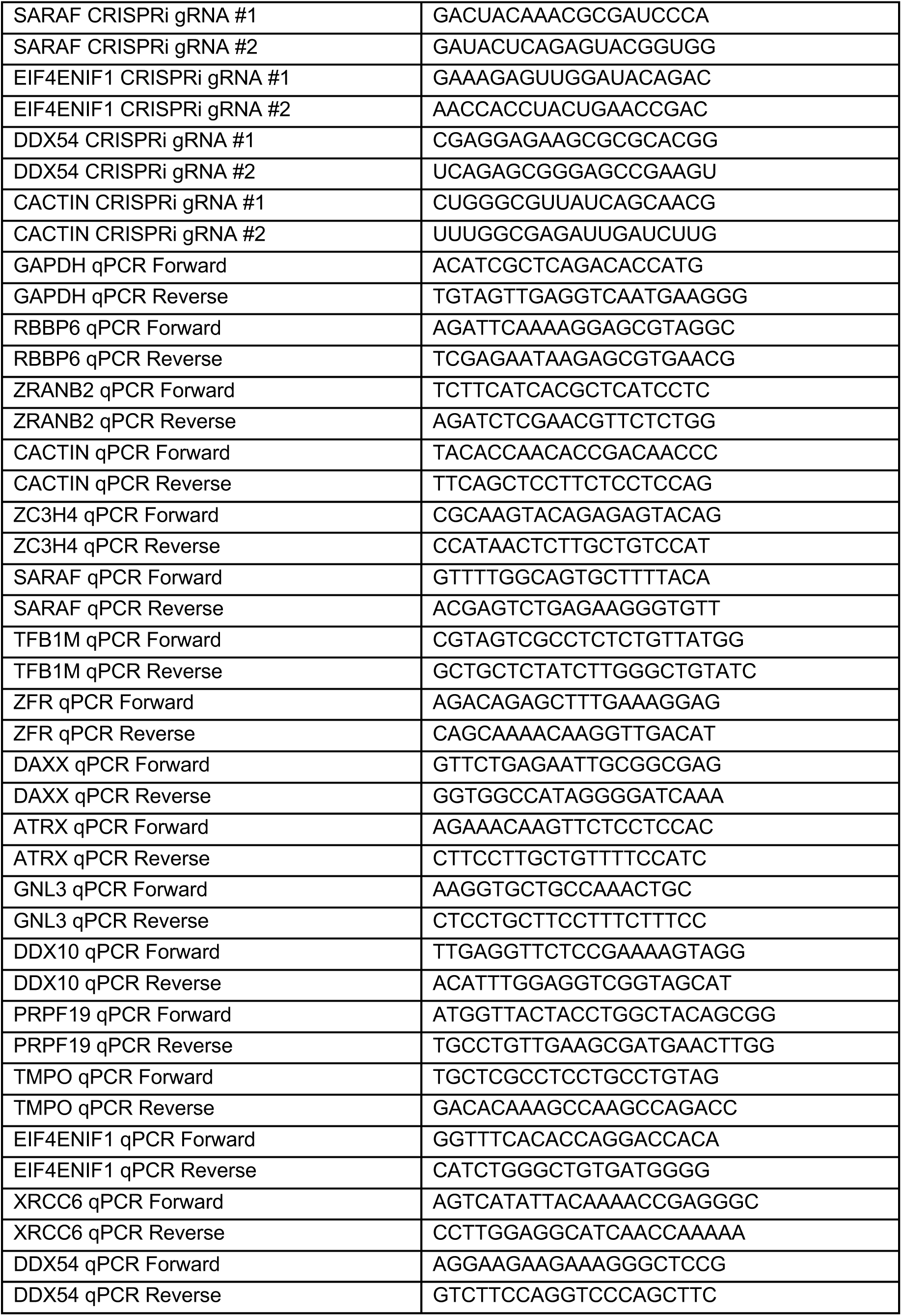
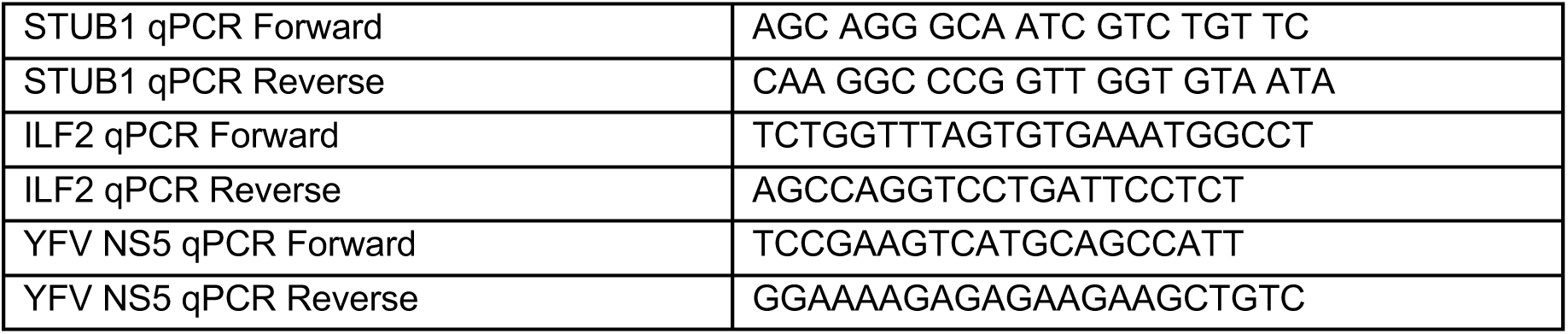
Oligonucleotides.

