## Supplementary Information for "Yellow Fever Virus Interactomes Reveal Common and Divergent Strategies of Replication and Evolution for Mosquito-borne Flaviviruses"

**Table S1. YFV-human proteomic data**

(A) Proteomic scoring
(B) Enrichment analysis
(C) Overlap statistics calculations

**Table S2. Targeted CRISPRi screen results.**

**Table S3. Evolutionary analysis of YFV interologs and RBBP6**

**Table S4. YFV-mosquito proteomic data**

(A) Proteomic scoring
(B) Ortholog mapping
(C) Enrichment analysis

**Table S5. Sequence and structural homolog analysis**

(A) FATCAT and NEEDLE analyses
(B) Foldseek forward search
(C) Foldseek reverse search
(D) Summary Statistics
(E) P values

**Table S6. Protein coding sequences used for YFV proteomics.**

**Table S7. List of antibodies used in this study**

**Table S8. List of oligonucleotides used in this study**

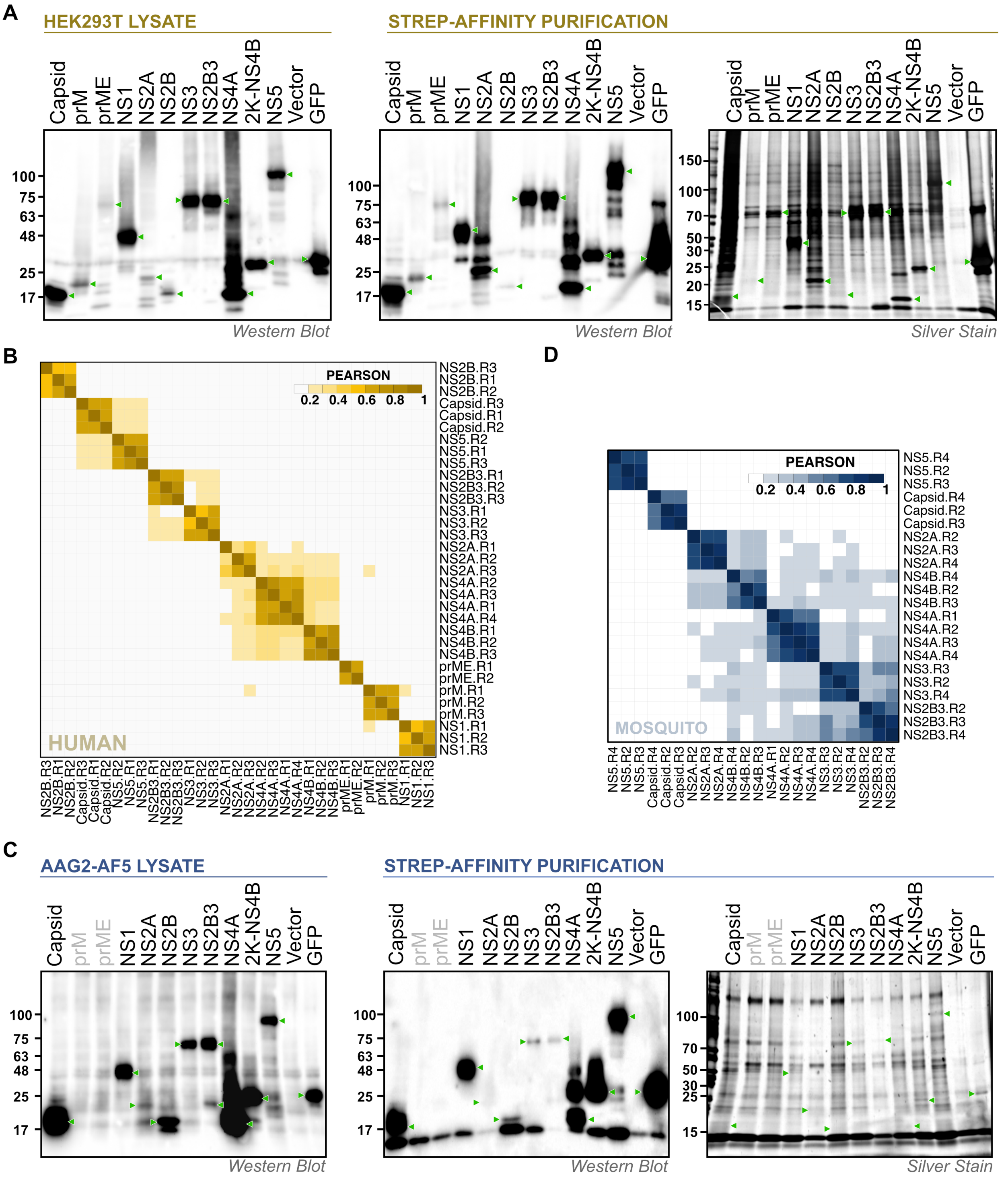


**Figure S1. YFV-host protein interaction quality control.**

(A) Western blotting and silver stain analysis of YFV protein expression and purification in human HEK 293T cells

(B) Hierarchical clustering of YFV-human proteomic data.

(C) Western blotting and silver stain analysis of YFV protein expression and purification in mosquito Aag2-AF5 cells
(D) Hierarchical clustering of YFV-mosquito proteomic data.

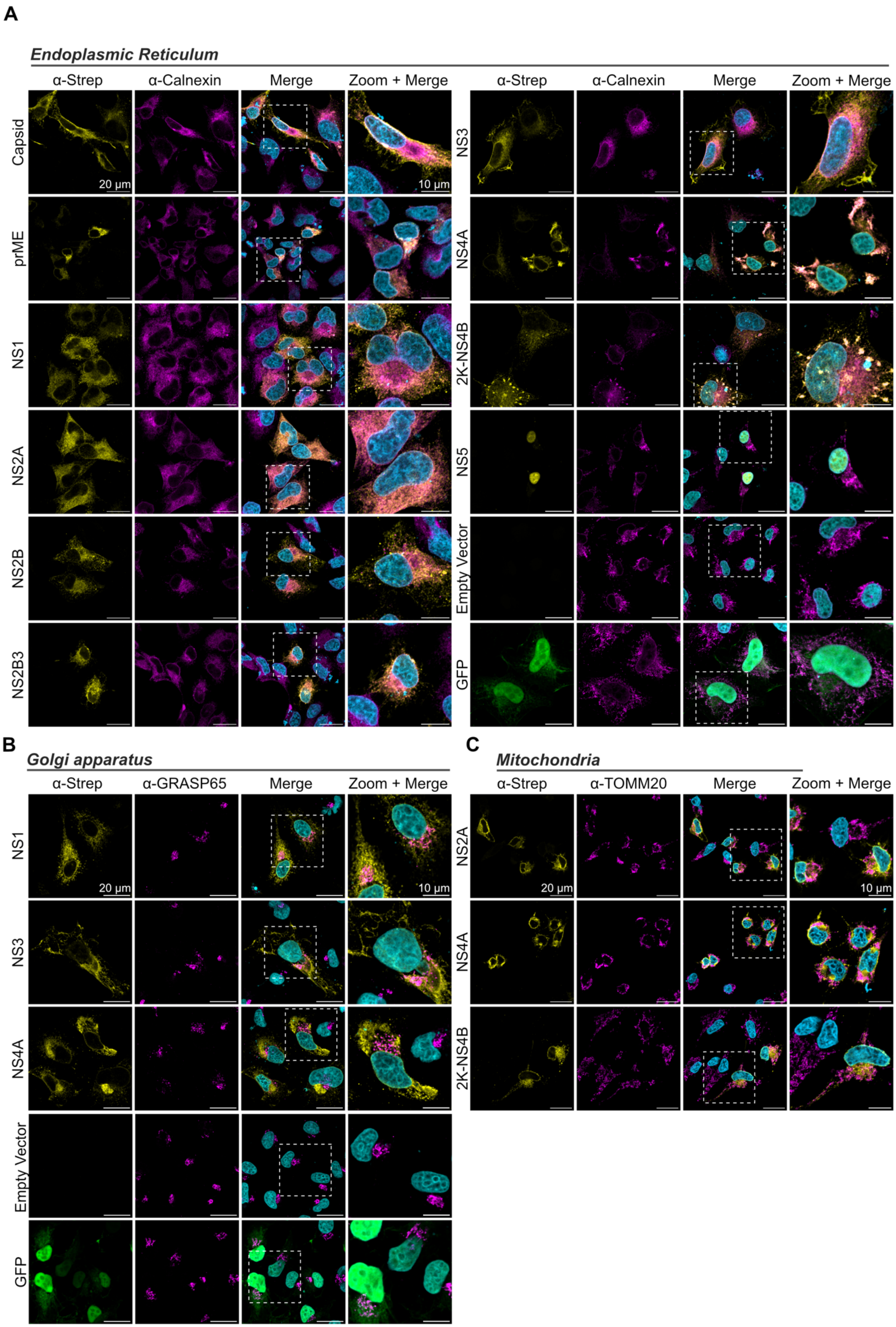
A
Endoplasmic Reticulum
α-Strep
α-Calnexin
Merge
Zoom + Merge
Capsid
prME
NS1
NS2A
NS2B
NS2B3
NS3
NS4A
2K-NS4B
NS5
Empty Vector
GFP
20 µm
10 µm
B
Golgi apparatus
α-Strep
α-GRASP65
Merge
Zoom + Merge
NS1
NS3
NS4A
Empty Vector
GFP
C
Mitochondria
α-Strep
α-TOMM20
Merge
Zoom + Merge
NS2A
NS4A
2K-NS4B

**Figure S2. Subcellular localization of YFV proteins in human cells.**

(A) ER co-staining (Calnexin)
(B) Golgi co-staining (GRASP65)
(C) Mitochondrial co-staining (TOMM20)

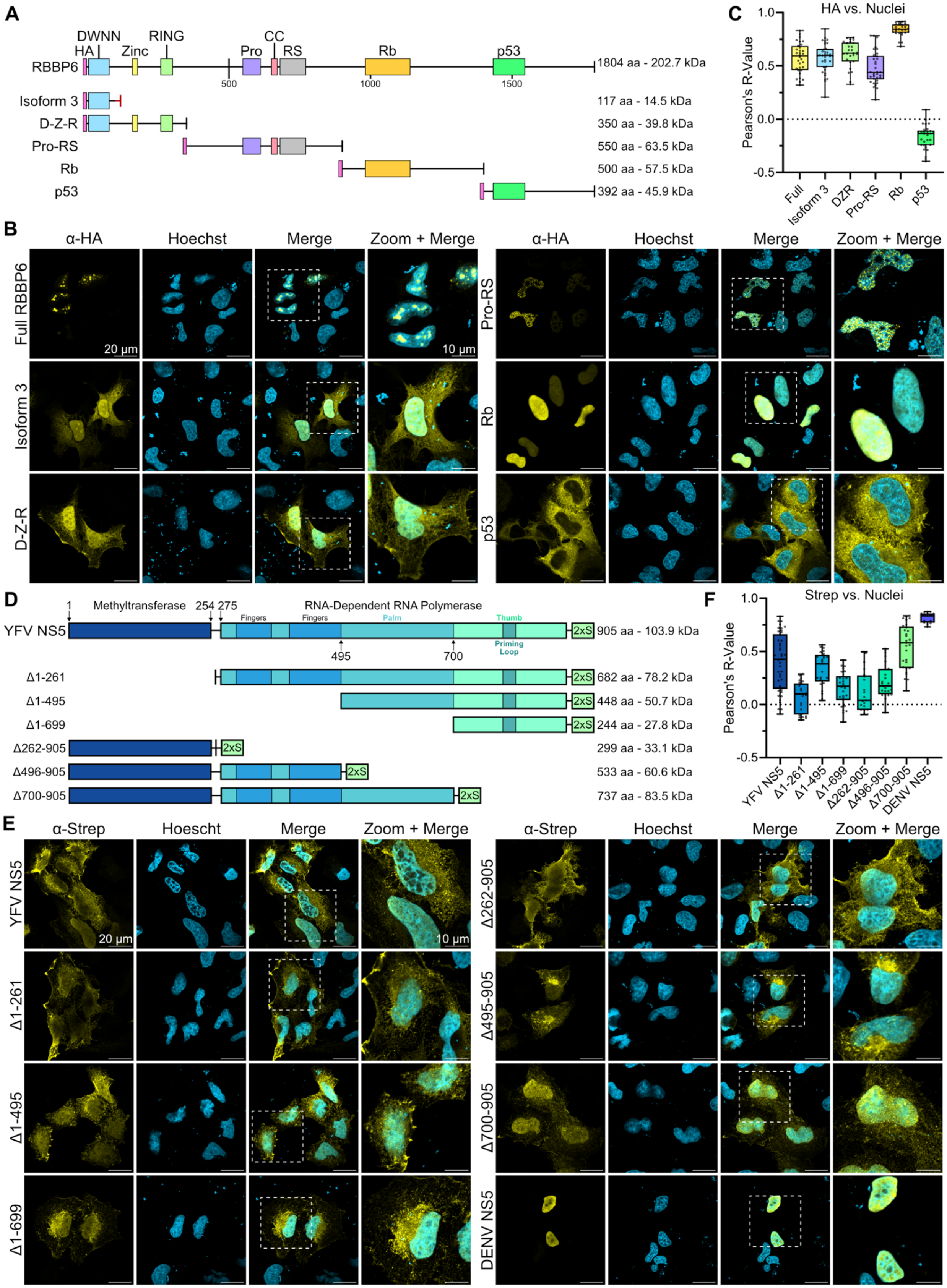

A
RBBP6
HA
DWNN
Zinc
RING
Pro
CC
RS
Rb
p53
1804 aa - 202.7 kDa
Isoform 3
117 aa - 14.5 kDa
D-Z-R
350 aa - 39.8 kDa
Pro-RS
550 aa - 63.5 kDa
Rb
500 aa - 57.5 kDa
p53
392 aa - 45.9 kDa
C
HA vs. Nuclei
Pearson's R-Value
Full
Isoform 3
DZR
Pro-RS
Rb
p53
B
α-HA
Hoechst
Merge
Zoom + Merge
Full RBBP6
Isoform 3
D-Z-R
Pro-RS
Rb
p53
20 μm
10 μm
D
YFV NS5
Methyltransferase
RNA-Dependent RNA Polymerase
Fingers
Palm
Thumb
Priming Loop
905 aa - 103.9 kDa
Δ1-261
682 aa - 78.2 kDa
Δ1-495
448 aa - 50.7 kDa
Δ1-699
244 aa - 27.8 kDa
Δ262-905
299 aa - 33.1 kDa
Δ496-905
533 aa - 60.6 kDa
Δ700-905
737 aa - 83.5 kDa
2xS
F
Strep vs. Nuclei
Pearson's R-Value
YFV NS5
Δ1-261
Δ1-495
Δ1-699
Δ262-905
Δ496-905
Δ700-905
DENV NS5
E
α-Strep
Hoescht
Hoechst
Merge
Zoom + Merge
YFV NS5
Δ1-261
Δ1-495
Δ1-699
Δ262-905
Δ495-905
Δ700-905
DENV NS5
20 μm
10 μm

**Figure S3. RBBP6 and YFV NS5 domain deletion subcellular localization.**

(A) Schematic of RBBP6 domain deletions
(B) Immunofluorescence microscopy of RBBP6 domain deletions
(C) RBBP6 domain deletion nuclear localization analysis using Pearson’s correlation with Hoescht
(D) Schematic of NS5 domain deletions
(E) Immunofluorescence microscopy of NS5 domain deletions
(F) NS5 domain deletion nuclear localization analysis using Pearson’s correlation with Hoescht

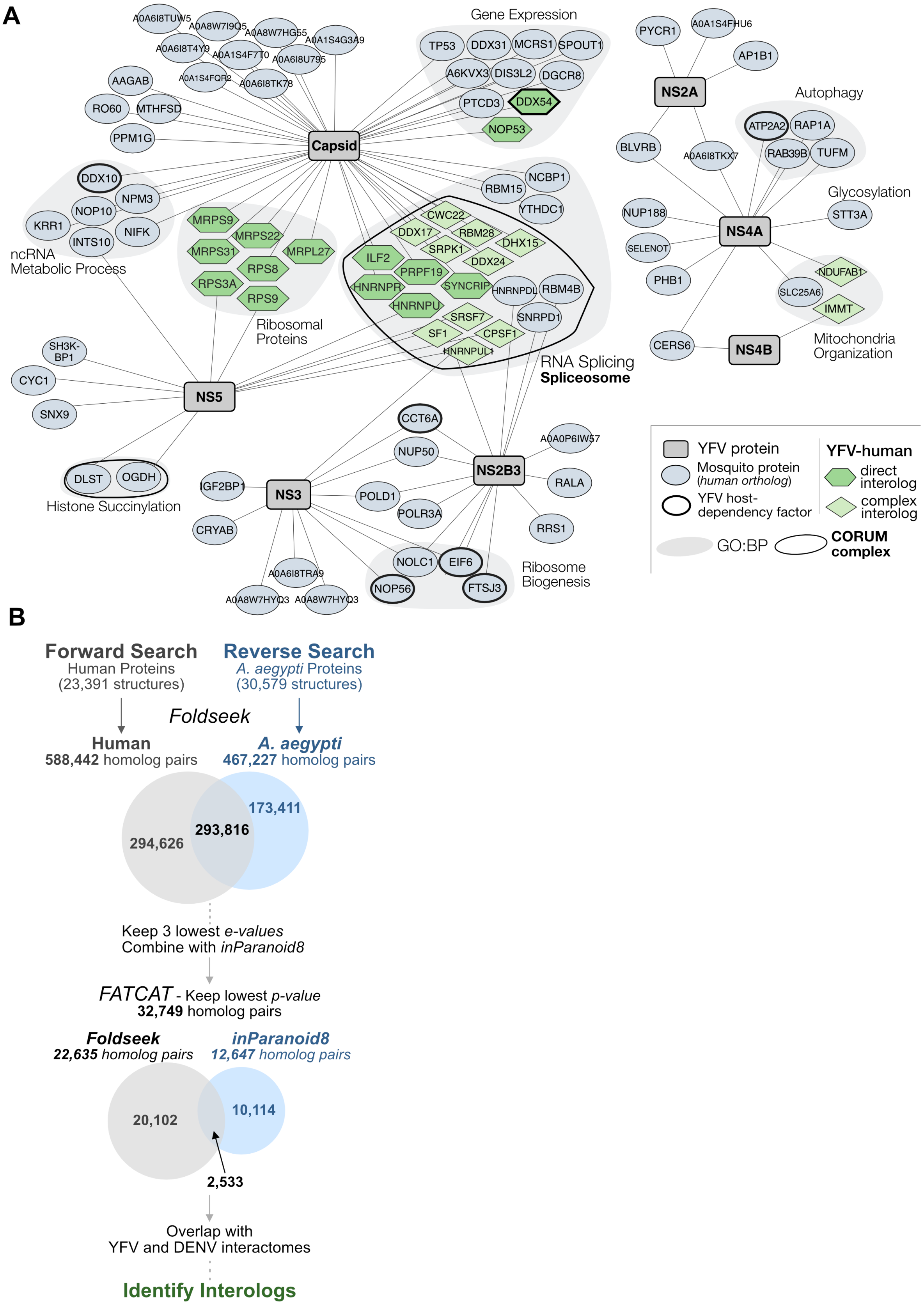

A
Gene Expression
TP53
DDX31
MCRS1
SPOUT1
A6KVX3
DIS3L2
DGCR8
PTCD3
DDX54
NOP53
A0A6I8TUW5
A0A8W7I9Q5
A0A8W7HG55
A0A1S4G3A9
A0A6I8T4Y9
A0A1S4F7T0
A0A6I8U795
A0A1S4FQR2
A0A6I8TK78
AAGAB
RO60
MTHFSD
PPM1G
Capsid
DDX10
NPM3
NOP10
KRR1
NIFK
INTS10
ncRNA
Metabolic Process
MRPS9
MRPS22
MRPS31
MRPL27
RPS8
RPS3A
RPS9
Ribosomal
Proteins
NCBP1
RBM15
YTHDC1
CWC22
DDX17
RBM28
DHX15
SRPK1
DDX24
ILF2
PRPF19
HNRNPR
SYNCRIP
HNRNPDL
RBM4B
HNRNPU
SRSF7
SNRPD1
SF1
CPSF1
HNRNPUL1
RNA Splicing
Spliceosome
PYCR1
A0A1S4FHU6
AP1B1
NS2A
Autophagy
ATP2A2
RAP1A
BLVRB
A0A6I8TKX7
RAB39B
TUFM
Glycosylation
NUP188
STT3A
NS4A
SELENOT
PHB1
NDUFAB1
SLC25A6
IMMT
CERS6
NS4B
Mitochondria
Organization
SH3K-
BP1
CYC1
SNX9
NS5
DLST
OGDH
Histone Succinylation
CCT6A
NUP50
A0A0P6IW57
NS2B3
RALA
IGF2BP1
NS3
POLD1
POLR3A
CRYAB
RRS1
NOLC1
EIF6
NOP56
FTSJ3
A0A6I8TRA9
A0A8W7HYQ3
A0A8W7HYQ3
Ribosome
Biogenesis
YFV protein
Mosquito protein
(human ortholog)
YFV host-
dependency factor
YFV-human
direct
interolog
complex
interolog
GO:BP
CORUM
complex
B
Forward Search
Human Proteins
(23,391 structures)
Reverse Search
A. aegypti Proteins
(30,579 structures)
Foldseek
Human
588,442 homolog pairs
A. aegypti
467,227 homolog pairs
294,626
293,816
173,411
Keep 3 lowest e-values
Combine with inParanoid8
FATCAT - Keep lowest p-value
32,749 homolog pairs
Foldseek
22,635 homolog pairs
inParanoid8
12,647 homolog pairs
20,102
10,114
2,533
Overlap with
YFV and DENV interactomes
Identify Interologs

**Figure S4. A YFV-mosquito HC-PPI network and structural interolog identification workflow**

(A) YFV-Mosquito HC-PPIs were identified after a tiered scoring approach using both MiST ($M > 0.67$ | $M > 0.60$ & protein complex member) and SAINT (score > 0.95 & BFDR ≤ 0.05). Viral baits (grey rectangles) are mapped to the mosquito proteins (blue grey ovals). Direct (protein level) and complex/pathway level evolutionary interologs are shown (green hexagons and diamonds). Human gene symbols are used when an appropriate evolutionary homolog is available (Table S4B). Proteins were grouped by GO:BP (grey underlay) and CORUM protein complex (black outline). Full scoring results are available in Table S4. Only ribosomal proteins that are also interologs are included for visual clarity.

(B) Schematic of the workflow used to identify the structurally similar protein pairs and interologs for both YFV and DENV.

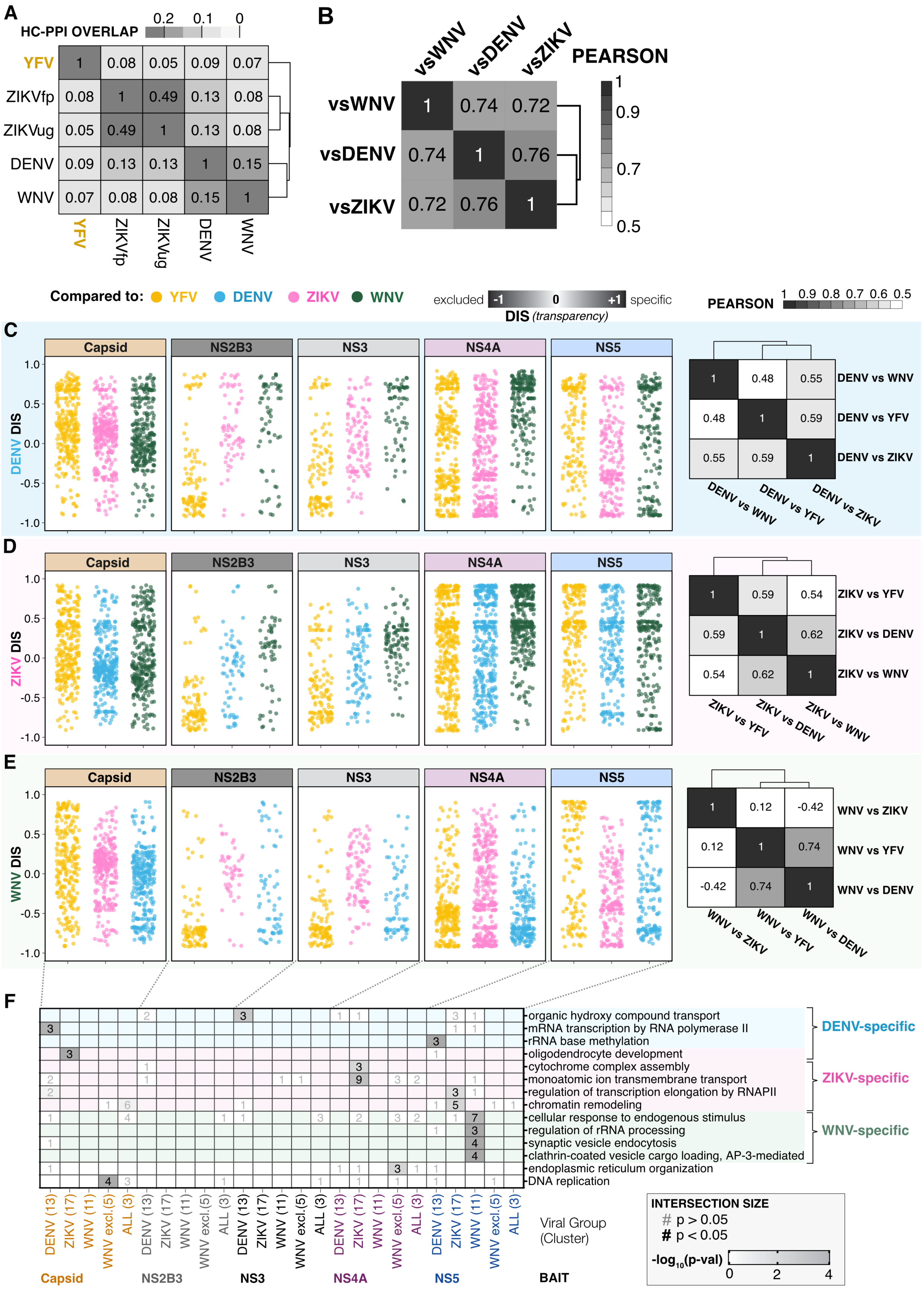

A
HC-PPI OVERLAP
0.2
0.1
0
YFV
ZIKVfp
ZIKVug
DENV
WNV
B
vsWNV
vsDENV
vsZIKV
PEARSON
Compared to:
YFV
DENV
ZIKV
WNV
excluded
specific
DIS (transparency)
C
Capsid
NS2B3
NS3
NS4A
NS5
DENV DIS
DENV vs WNV
DENV vs YFV
DENV vs ZIKV
D
ZIKV DIS
ZIKV vs YFV
ZIKV vs DENV
ZIKV vs WNV
E
WNV DIS
WNV vs ZIKV
WNV vs YFV
WNV vs DENV
F
organic hydroxy compound transport
mRNA transcription by RNA polymerase II
rRNA base methylation
oligodendrocyte development
cytochrome complex assembly
monoatomic ion transmembrane transport
regulation of transcription elongation by RNAPII
chromatin remodeling
cellular response to endogenous stimulus
regulation of rRNA processing
synaptic vesicle endocytosis
clathrin-coated vesicle cargo loading, AP-3-mediated
endoplasmic reticulum organization
DNA replication
DENV-specific
ZIKV-specific
WNV-specific
DENV (13)
ZIKV (17)
WNV (11)
WNV excl.(5)
ALL (3)
Viral Group (Cluster)
Capsid
NS2B3
NS3
NS4A
NS5
BAIT
INTERSECTION SIZE
# p > 0.05
# p < 0.05
-log10(p-val)

**Figure S5. Differential interaction score (DIS) across flaviviruses.**

(A) Heatmap of percent overlap of HC-PPIs across the indicated flavivirus datasets.

(B) Heatmap of Pearson's correlations of the DIS distributions for YFV versus the indicated flavivirus datasets.

(C-E) Plot of DENV, ZIKV, and WNV DIS distributions, separated by the viral bait. Color indicates virus used for DIS comparison. Heatmap of Pearson's correlations of the DIS distributions for indicated flavivirus versus the other flaviviruses.

(F) Heatmap showing the degree of enrichment ($\log_{10}$p-value, yellow scale bar) for unique terms in clusters associated with DENV-, ZIKV-, and WNV-specific interactions, grouped by viral bait. In each tile, the number of interactions associated with that enrichment term is bolded if it was significant ($p < 0.05$).

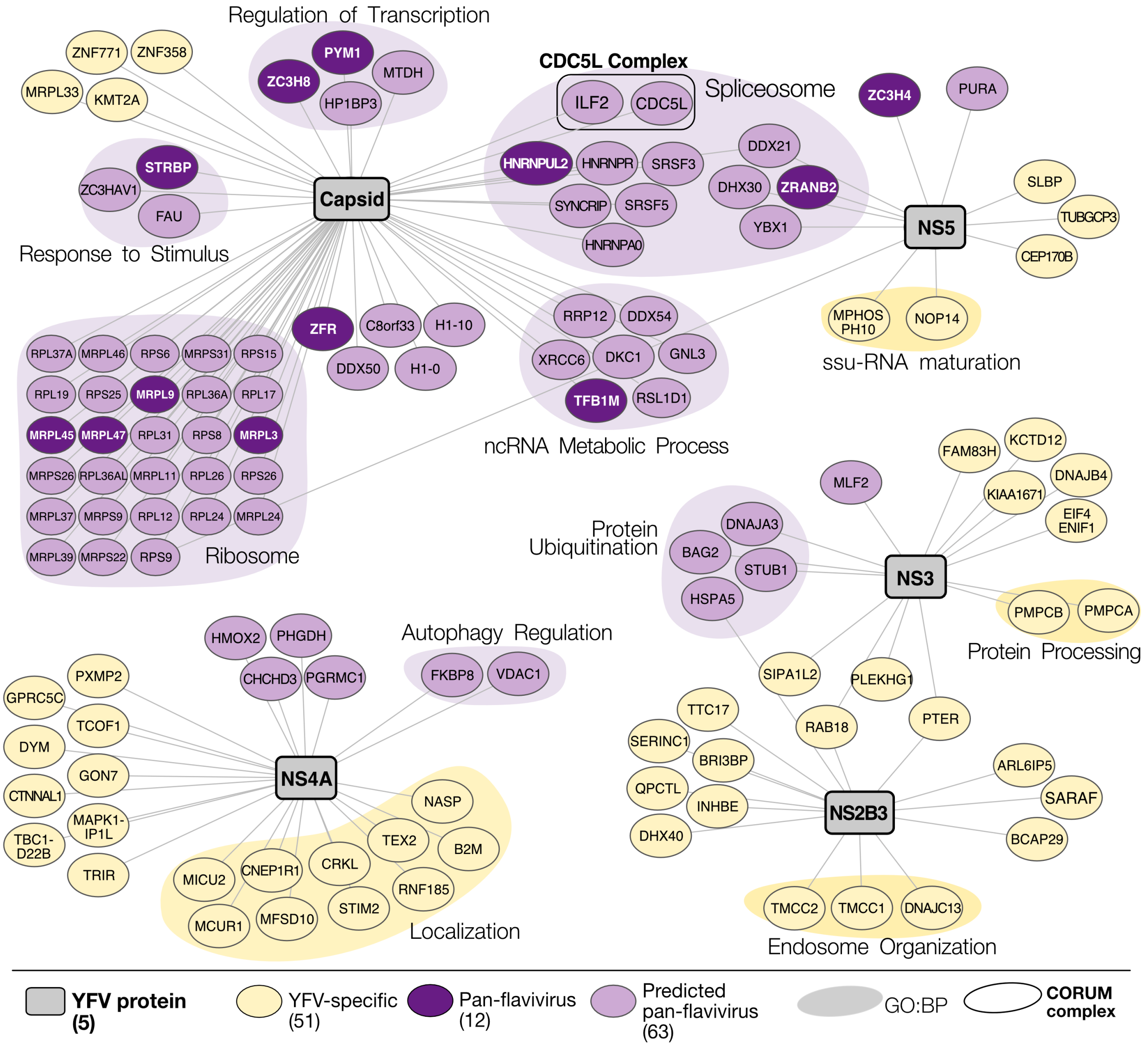


**Figure S6. A YFV-human HC-PPI network with pan-flavivirus and YFV-specific interactions.**

YFV-Human HC-PPIs were annotated for pan-flavivirus and YFV-specific interactions based on predictions from cluster 3 and 1 Figure 5B, respectively. Viral baits (grey) are mapped to the human proteins (yellow or purple). GO:BP enrichment and CORUM complexes are also annotated.

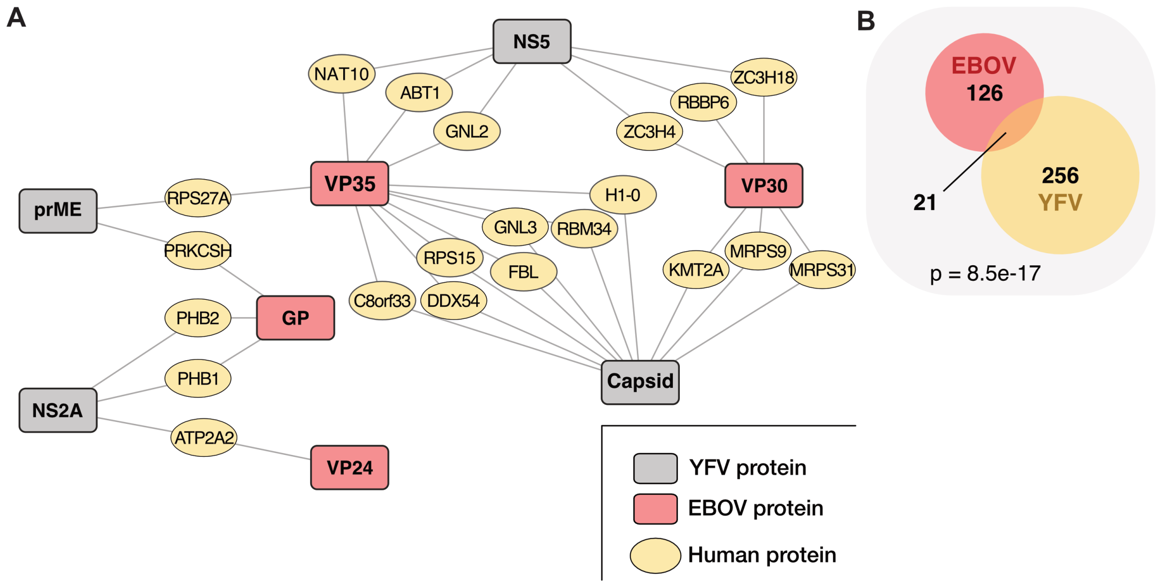


**Figure S7. Overlap of YFV and EBOV interactomes.**

(A) Overlap of YFV and EBOV interactomes
(B) Venn diagram and Fisher's exact test of overlap in (A)
